# Role of Myc family proteins in transcriptional regulation of growth and oncogenic transformation in fusion-positive rhabdomyosarcoma

**DOI:** 10.64898/2025.12.01.690506

**Authors:** Bishwanath Chatterjee, Pawan K. Raut, Hana Kim, Rachel A. Hoffman, Puspa R. Pandey, Shabir Zargar, Wenyue Sun, Benjamin Z. Stanton, Frederic G. Barr

**Affiliations:** Laboratory of Pathology, Center for Cancer Research, National Cancer Institute, Bethesda, MD 20892-1500; Center for Childhood Cancer Research, Abigail Wexner Research Institute at Nationwide Children’s, Columbus, OH 43205

**Author notes:** B. Chatterjee, P. K. Raut and H. Kim contributed equally to this article. Corresponding author: Frederic G. Barr, Laboratory of Pathology, Center for Cancer Research, National Cancer Institute, 10 Center Drive, Room 2S235D, Bethesda, MD 20892-1500. **Conflict of interest:** The authors declare no potential conflicts of interest. **Statement of Significance:** MYCN and/or MYC expression contributes to growth and oncogenic transformation in fusion-positive rhabdomyosarcoma cells and engineered myoblast models by facilitating PAX3::FOXO1 activation of *FGF8* and increasing expression of Myc-specific pathways.

**Keywords:** MYC, MYCN, FGF8, fusion gene, oncogenic transformation

## Abstract

Fusion-positive rhabdomyosarcoma (FP-RMS) is driven by a PAX*3::FOXO1 (P3F)* or *PAX7::FOXO1* fusion gene. After finding that MYCN is required for P3F-induced oncogenic transformation in a human myoblast model of FP-RMS, we further investigated the role of Myc family proteins in FP-RMS. Expression studies revealed that myoblast models and a subset of FP-RMS lines have predominant MYCN or MYC expression whereas other FP-RMS lines have both high MYCN and MYC expression. In myoblast models, MYCN was required for optimal P3F binding to and high-level activation of *FGF8*, a P3F target that is necessary and sufficient for oncogenic transformation. *MYCN* or *MYC* knockdown suppressed transformation in myoblast and FP-RMS lines with dominant MYCN or MYC expression, respectively. In FP-RMS lines with high MYCN and MYC expression, there was partial loss of transformation when one gene was targeted and complete loss when both genes were targeted. Despite the loss of oncogenic activity in lines with knockdown of a dominant Myc family member, *FGF8* expression was not decreased. Transcriptomic analyses revealed that P3F target genes were not affected by *MYCN* or *MYC* knockdown, and instead a group of Myc-specific targets was down-regulated. Collectively, these results indicate that MYCN or MYC is functionally dominant in myoblast models and FP-RMS lines with dominant expression of one Myc family protein whereas MYCN and MYC are functionally redundant in FP-RMS cell lines with high expression of both. These Myc family proteins contribute to oncogenic properties by facilitating P3F activation of *FGF8* and increasing expression of Myc-specific targets.

## Introduction

Rhabdomyosarcoma (RMS) represents a family of soft tissue sarcomas which are associated with the skeletal muscle lineage and occur in the pediatric population [1]. RMS was originally classified into two major subtypes, alveolar (ARMS) and embryonal (ERMS), based on histological features. Following the discovery of fusion genes in most ARMS tumors, RMS can now be divided into fusion-positive (FP-RMS) and fusion-negative (FN-RMS) categories [2]. FP-RMS is characterized by a recurrent 2;13 or 1;13 chromosomal translocation, which breaks within the *PAX3* or *PAX7* and *FOXO1* genes to generate a *PAX3::FOXO1* (*P3F*) or *PAX7::FOXO1* (*P7F*) fusion gene [3, 4]. These fusion genes encode fusion transcription factors that are oncogenic drivers in FP-RMS [2].

Myc family members, including MYC, MYCN and MYCL, are transcription factors containing basic-helix-loop-helix and leucine zipper domains [5]. These Myc proteins are often overexpressed or activated in different types of cancers and contribute to the growth and oncogenic properties of these cancers [6]. Genome-wide screens identified a subset of FP-RMS tumors with amplicons derived from the 2p24 chromosomal region where the *MYCN* locus is situated [7]. Focused analysis of the *MYCN* gene revealed FP-RMS cases with increased *MYCN* copy number and increased *MYCN* expression at both the RNA and protein levels [8–10]. In addition to *MYCN* amplification in FP-RMS cases, *MYCN* is a downstream transcriptional target of P3F and is thus generally upregulated in FP-RMS tumors [9, 11].

In previous studies, we found that P3F expressed from either a constitutive or doxycycline-inducible construct cannot induce transformation in immortalized human Dbt myoblasts [12, 13]. Of note, endogenous *MYCN* is not highly induced by P3F in these myoblasts. While high-level constitutive MYCN expression also cannot transform these myoblasts, the combination of constitutive MYCN and either constitutive or inducible P3F does transform these myoblasts. Furthermore, Dbt cells expressing P3F and MYCN readily form tumors when injected intramuscularly in immunodeficient mice. In contrast, Dbt cells with only P3F expression develop tumors at a much slower rate and Dbt cells with only MYCN expression do not form tumors [13]. These findings indicate that MYCN collaborates with P3F to transform the myoblasts in vitro and enhance tumor formation in vivo.

In the current study, we investigated the role of Myc family proteins, particularly MYC and MYCN, in FP-RMS. We performed a comprehensive comparison of expression of these Myc family members at the RNA and protein levels in the human myoblast model system and in FP-RMS cell lines derived from human tumors. We used gene expression profiling and chromatin immunoprecipitation studies to investigate downstream changes associated with expression of Myc family proteins in the myoblast model system. Finally, we performed knockdown experiments to dissect the dependence on MYC and MYCN in growth and transformation in the myoblast system and FP-RMS lines.

## Materials and Methods

### Cell culture

The engineered immortalized human Duchenne muscular dystrophy myoblast cell line (Dbt) and human RMS cell lines were cultured as described previously [12].The sources of the cell lines are as follows: Dbt - D. Trono; RH30 - ATCC; RH28 - B. Emanuel; RH5 -J. Khan; CW9019 - J. Biegel; RH41 - C. Linardic; MP4 - T. Cripe; NCI-RMS-097 - L. Helman; RH10 - P. Houghton. Verification of cell line identity was performed by short tandem repeat genotyping with the AmpFISTR Profiler Plus PCR amplification kit (Applied Biosystems). Cell lines were periodically checked with a PCR-based Mycoplasma detection kit (ATCC, # 30-1012K) to rule out the possibility of Mycoplasma contamination.

### Transfection and lentiviral transduction

Dbt cells were transduced either with a constitutive MYCN expression construct or empty vector and with an inducible P3F expression construct or empty vector. The resulting lines expressing MYCN only, P3F only or both MYCN and P3F are designated as Dbt-MYCN, Dbt-iP3F and Dbt-MYCN-iP3F, respectively. The pCMV6-Entry vector expressing FGF8b cDNA or empty vector (EV; Origene) was stably transfected using Lipofectamine 3000 (ThermoFisher Scientific). CRISPR/Cas9 targeting of *FGF8*, *MYC*, *MYCN*, *P3F* and *P7F* was performed with a single guide RNA (designated as CR gRNA-1) (Supp Table 1) cloned into lentiCRISPR v2 (GenScript) and compared with a control nontargeting vector lentiCRISPR v2 (CR-Control). For combined targeting of both *MYC* and *MYCN*, two guide RNAs (designated as CR gRNA-1 and CR gRNA-2) were used for each gene. Lentiviral particles were packaged using HEK293T cells and human immunodeficiency virus-based pPACK-H1 Packaging Plasmid Mix (System Biosciences).

### RNA extraction and gene expression profiling

Total RNA was extracted using the RNAeasy Kit (Qiagen). Taqman gene expression assays (Life Technologies) were performed using cDNA prepared by Superscript IV to quantify expression of *P3F* (Hs03024825_ft), *FGF8* (Hs00171832_m1), *MYC* (Hs00153408_m1), *MYCN* (Hs00232074_m1) and *GAPDH* (Hs02758991) (ThermoFisher). The 2–ΔΔCt method [14] was used to quantify the test gene expression change relative to the GAPDH control.

The RNA-Seq datasets were processed using the NIH CCBR pipeline as previously described [15]. Briefly, sequence quality was checked with FastQC (version 0.11.9). Adapter sequences were removed using Cutadapt [16] before alignment to the hg38 reference genome using STAR v2.7.6a [17] in two-pass mode. Gene expression levels were quantified using RSEM [18]. Downstream analysis and visualization were performed within the NIH Integrated Analysis Platform (NIDAP) using R programs developed by a team of NCI bioinformaticians on the Foundry platform (Palantir Technologies). The raw counts matrix was imported into the NIDAP platform, where genes were filtered for low counts (<1 cpm) and normalized by quantile normalization using the limma package [19]. Differentially expressed genes were calculated using limma-Voom [20]. Genes with p- values <= 0.05 and fold change of >= |1.2| were considered significantly differentially expressed. Pathway enrichment analysis was performed using Fisher’s Exact Test (L2P Package on GitHub: https://github.com/ccbr/l2p) and bubble plots represent selected significantly enriched pathways. The data presented in this article have been deposited in and are available from the dbGaP database under accession phs004312.v1.p1.

### Protein extraction and western blotting

Protein extraction and western blotting were performed as described previously [21]. Membranes were incubated overnight with antibodies against FOXO1 (1:1,000, catalog no. 2880; Cell Signaling Technology), FGF8 (1:1,000, catalog no. 16124, Sino Biological), MYCN (B8.4.B) (1:1,000, catalog no. sc-53993, Santa Cruz), MYC (Y69) (1:1,000, catalog no. ab32072, Abcam) and GAPDH (1:2,000, catalog no. sc-25778; Santa Cruz Biotechnology).

### Proliferation, focus formation and clonogenic assays

To evaluate population growth, cells were seeded at a density of 1,000-1,500 cells per well in 96-well plates, and cell confluency was monitored on the IncuCyte S3 Live-Cell Analysis System (Essen BioScience) every 6 hrs for 90 hrs, as described previously [22]. Data were evaluated using IncuCyte^®^ Basic Analysis Software (2023A Rev2 GUI).

For Dbt derivatives, the focus formation assay (for assessing *in vitro* oncogenic transformation) and the clonogenic assay (for assessing colony growth on plastic plates) were performed as previously described in 60 mm dishes containing a 50:50 mixture of F-10 and DMEM [23]. Medium with doxycycline (500 ng/ml) was changed every 3 days during the 12-16 days assay period. For focus formation assays in FP-RMS cell lines, 360-720 FP-RMS cells and 360,000 NIH3T3 fibroblasts were co-cultured into 60 mm dishes in growth medium. For the clonogenic assay, 360-720 FP-RMS cells were cultured into 60 mm dishes in a 50:50 mixture of conditioned medium and selected growth medium. The medium was changed every 3-4 days during the 14-18 days assay period for both clonogenic and focus formation assays. At the end of the assay period, the cells were fixed with methanol, stained with Giemsa stain (Sigma-Aldrich) and imaged with ChemiDoc XRS+ Imaging System (Bio-Rad). For each condition, foci or colonies were counted using Image J software [24].

### Chromatin immunoprecipitation (ChIP) studies of P3F binding

ChIP was performed in two biological replicates as previously described [25]. Reads were aligned to hg38 and processed using the ENCODE ChIP-seq pipeline, version 2.1.2 [26]. The WashU Epigenome Browser was used for data visualization [27]. Quantification of ChIP signal relative to input was obtained from the fold-change bigwig files produced by the ENCODE pipeline, and the maximum fold-change within each P3F binding peak was determined using the Genomic Ranges package in R [28]. Data were plotted using the ggplot2 package in R [29].

Statistical analysis

Cell culture experiments including clonogenic and focus formation assays were performed with 3 biological replicates unless otherwise stated, while population growth studies were carried out with 6 biological replicates for each condition. Representative results are shown in the figures and are presented as mean ± SE. Quantitative data were analyzed for significance using a two-sided unpaired Student *t* test to compare CR-Control with different knockdown groups including CR-P3F/P7F, CR-MYC, CR-MYCN and CR-MYC-MYCN. Results were deemed to be statistically significant when P ≤ 0.05. Prism 8.4.3 software (GraphPad Software) was used for data presentation and statistical analysis.

## Results

### Myc family gene expression in engineered Dbt myoblast and FP-RMS lines

Based on initial evidence that MYCN contributes to oncogenicity in FP-RMS, we measured expression of Myc family members (*MYCN*, *MYC* and *MYCL)* in our engineered Dbt myoblasts. We used both RNA-Seq and quantitative RT-PCR (qRT-PCR) to analyze RNA expression in parental (Pa) and tumor-derived (TD) cells from Dbt-MYCN-iP3F and Dbt-iP3F myoblasts. Our analysis showed that the Dbt-iP3F-Pa lines had moderate levels of *MYC* and low levels of *MYCN* in the presence or absence of P3F (Fig. 1A, Fig. S1A-B). Although the *MYC* level did not significantly change in Dbt-iP3F-TD cells, we observed a 10-fold increase in *MYCN* expression (3.4 log2 Fold Change [FC], p=0.00054) in Dbt-iP3F-TD lines compared to Dbt-iP3F-Pa lines. As expected, engineered *MYCN* expression resulted in very high *MYCN* levels in both Dbt-MYCN-iP3F-Pa and Dbt-MYCN-iP3F-TD lines. Along with this increase in *MYCN* expression, there is a 9-fold decrease in *MYC* expression (3.2 log2 FC, p<0.00001) in Dbt-MYCN-iP3F-Pa and Dbt-MYCN-iP3F-TD lines compared to the Dbt-iP3F-Pa and Dbt-iP3F-TD lines. Lastly, the expression of *MYCL* did not change as much as *MYC* or *MYCN* in these cell lines. *MYCL* expression was slightly up-regulated (2-3 fold) by P3F induction in the Dbt-iP3F-Pa (1.1 log2 FC, p=0.0077) and Dbt-MYCN-iP3F-Pa (1.5 log2 FC, p=0.027) lines, and overall *MYCL* levels did not change in TD lines compared to uninduced parental cells.

**Figure 1.**
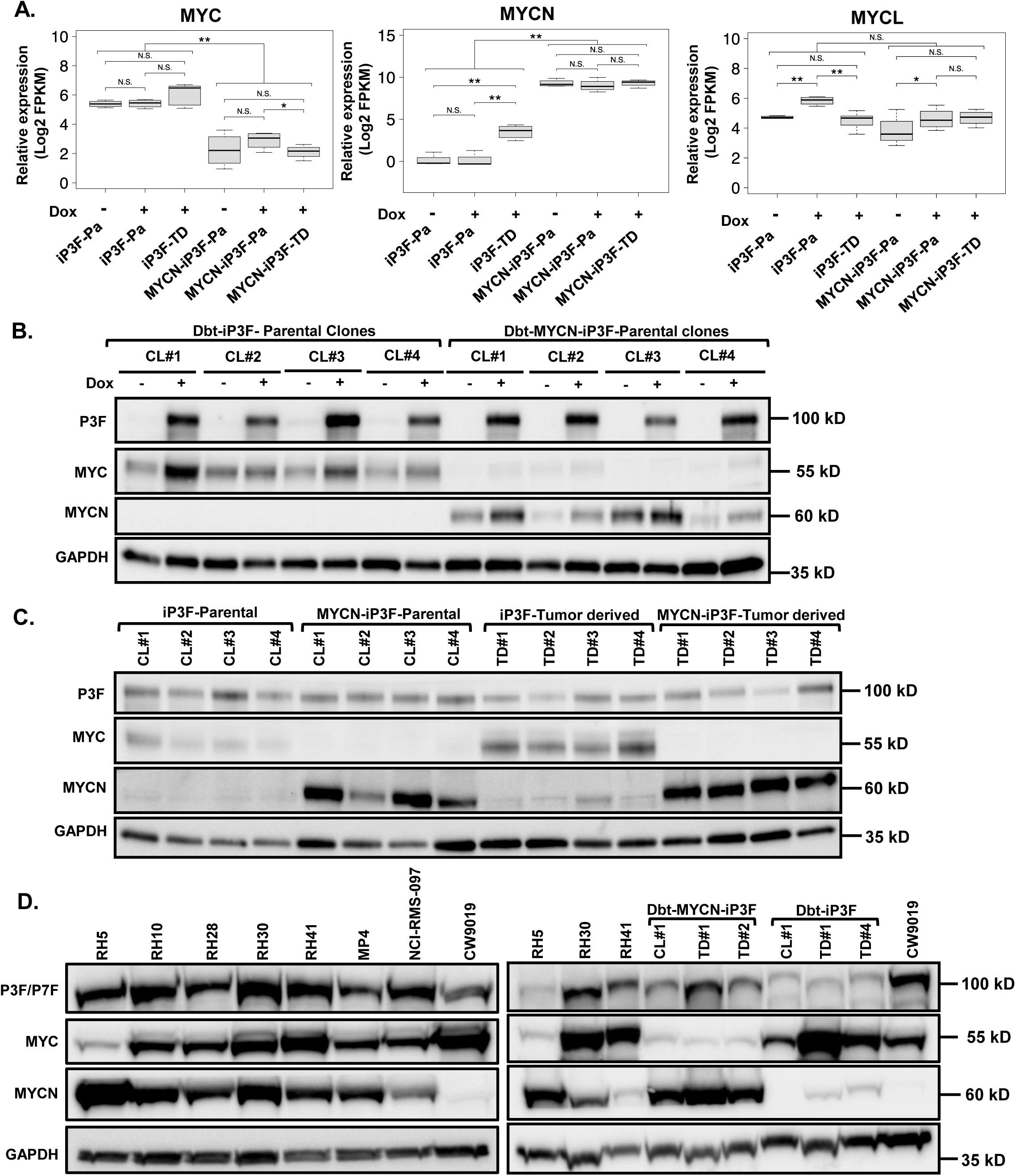
Expression of Myc family members in engineered myoblasts and fusion-positive rhabdomyosarcoma (FP-RMS) cell lines. **A.** RNA-Seq analysis of *MYC*, *MYCN* and *MYCL* RNA expression in parental (Pa) and tumor-derived (TD) myoblast lines with (+) or without (-) 500 ng/ml doxycycline (Dox) induction of PAX3::FOXO1 (P3F) expression. Each measurement is expressed as mean ± SE of 4 biological replicates (subclones). Significance levels are: *, *P <* 0.05; **, *P <* 0.01; ***, *P* < 0.001; N.S., not significant. **B.** P3F, MYCN and MYC protein expression in parental myoblast subclones (CL) with (+) or without (-) doxycycline induction of P3F. **C.** P3F, MYCN and MYC protein expression in parental and TD myoblast subclones with doxycycline induction of P3F. **D.** P3F/P7F, MYCN and MYC protein expression in engineered myoblasts and FP-RMS lines. Left, FP-RMS cell lines. Right, engineered myoblast lines and FP-RMS lines.

At the protein level, higher MYCN levels were seen in Dbt-MYCN-iP3F lines (both Pa and TD) compared to Dbt-iP3F lines while higher MYC levels were seen in Dbt-iP3F (both Pa and TD) compared to Dbt-MYCN-iP3F lines (Fig. 1B-C); these expression patterns are similar to the RNA findings. In addition, MYCN protein levels increased in Dbt-iP3F-TD lines compared to Dbt-iP3F-Pa lines in agreement with the RNA studies (Fig. 1C). In contrast to the RNA results, MYC protein expression increased in Dbt-iP3F-Pa lines after P3F induction. Furthermore, there was an increase in the MYC protein level in Dbt-iP3F-TD lines compared to Dbt-iP3F-Pa lines (with P3F induction), which is substantially more than the change seen at the RNA level. Based on the predominant Myc family protein, we hypothesize that the oncogenic changes in the Dbt-MYCN-iP3F-TD cells are associated with high MYCN expression while the oncogenic changes in the Dbt-iP3F-TD cells are associated with high MYC expression.

Next, we determined *MYCN* and *MYC* expression in native FP-RMS lines (Fig. 1D and Fig. S1C-E). RNA and protein analyses demonstrated varying amounts of MYC and MYCN among the lines. Although there is an overall gradient of MYC and MYCN expression, three groups can be discerned: MYC/MYCN co-dominant, MYC-dominant and MYCN-dominant. The MYC/MYCN co-dominant group (exemplified by RH28 and RH30) express substantial amounts of both MYC and MYCN. The MYC-dominant group (exemplified by CW9019) shows a high level of MYC and a low level of MYCN whereas the MYCN-dominant group (exemplified by RH5) expresses a high level of MYCN and a low level of MYC. Copy number analysis of RH5 cells demonstrated *MYCN* amplification (data not shown) that results in 5-10-fold higher *MYCN* RNA levels than the other lines and a corresponding decrease in *MYC* expression. The MYC expression in MYC-dominant CW9019 cells is similar to the MYC level in MYC/MYCN co-dominant cells such as RH30 and thus the mechanism for the very low level of MYCN expression in CW9019 is not known. Although CW9019 is the only *P7F*-positive RMS line in this panel, *MYCN* is amplified in another *P7F*-positive cell line [30] and is highly expressed in some *P7F*-positive tumors (data not shown) and thus the low MYCN expression in CW9019 is not specifically associated with *P7F* fusion status.

We next compared the level of MYC and MYCN expression in these FP-RMS lines and the engineered myoblast lines. These results revealed similarities in MYC and MYCN expression levels in the native FP-RMS lines and our myoblast models of FP-RMS (Fig. 1D). In particular, we find that the MYCN-dominant RH5 and Dbt-MYCN-iP3F-TD lines have similar high MYCN and low MYC levels. In addition, the MYC-dominant CW9019 and Dbt-iP3F-TD lines express similarly high MYC and low MYCN levels. These findings support that our myoblast system provides a useful model of MYCN-dominant and MYC-dominant FP-RMS.

### Dependency of some P3F target genes on Myc family protein expression

To elucidate P3F targets that contribute to oncogenic transformation of the engineered myoblast lines, we performed a transcriptomic analysis focusing on P3F target genes [31]. We hypothesize that some P3F targets needed for oncogenic transformation depend on the presence of Myc family proteins for transcriptional activation. We thus looked for P3F target genes that are upregulated by doxycycline in the Dbt-MYCN-iP3F-Pa, Dbt-MYCN-iP3F-TD and Dbt-iP3F-TD lines (all of which highly express a Myc family protein and are transformed) but not in the Dbt-iP3F-Pa lines (that do not highly express a Myc family protein and are not transformed). These differential expression studies indicated that 22 out of 1,010 P3F target genes were up-regulated and 6 were down-regulated in Dbt lines with P3F and high Myc family expression and not in lines without P3F and/or high Myc family expression (cutoff of greater than 2-fold change, p<0.05) (Fig. 2A and data not shown). Among the top ten up-regulated P3F targets, *FGF8* showed the highest average expression among TD lines (80-fold change, p<0.00001) and thus *FGF8* fits the criteria for a highly expressed Myc family-dependent P3F target gene (Fig. 2A).

**Figure 2.**
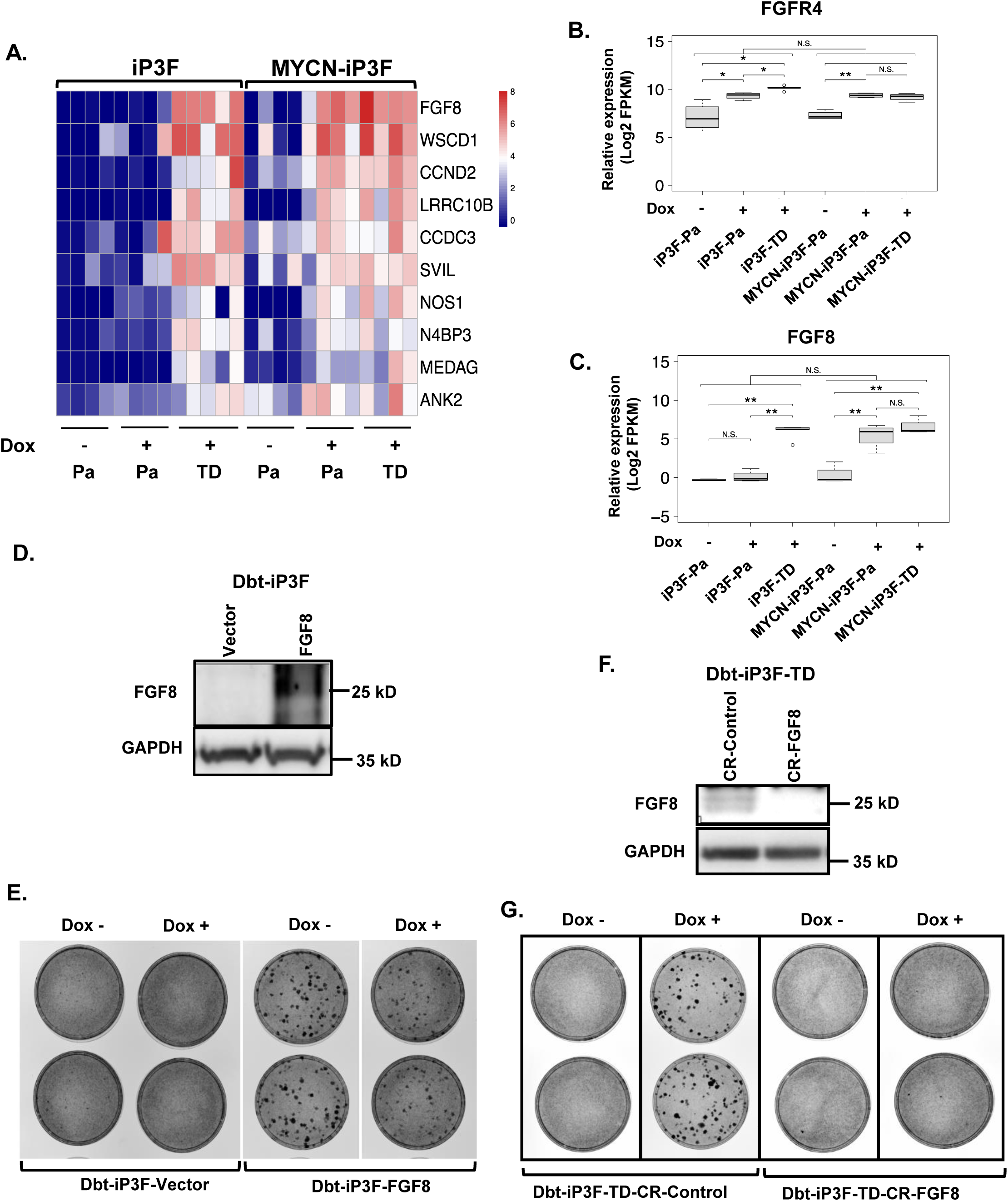
Expression and function of P3F target genes in engineered Dbt cell lines. **A.** Heat map of RNA-Seq data for the most highly upregulated P3F target genes in parental (Pa) and tumor-derived (TD) myoblast lines with (+) or without (-) 500 ng/ml doxycycline (Dox) induction of P3F expression. **B, C.** RNA-Seq analysis of *FGFR4* (B) and *FGF8* (C) RNA expression in myoblast lines with (+) or without (-) Dox induction of P3F expression. Each measurement is expressed as mean ± SE of 3-4 biological replicates (subclones). Significance levels are described in Fig. 1. **D, E.** Western blot analysis of FGF8 protein expression (D) and focus formation assay (E) of Dbt-iP3F-Pa cells transfected with control or FGF8 expression construct. **F, G.** Western blot analysis of FGF8 protein expression (F) and focus formation assay (G) of Dbt-iP3F-TD line transduced with control or FGF8-specific knockdown construct.

Next, we used RNA-Seq and qRT-PCR to measure expression of selected P3F target genes [31] in the parental and TD myoblast lines with or without P3F stimulation. Our analysis confirmed two categories of target genes. In one set, exemplified by *FGFR4*, the target genes are stimulated at similar levels by P3F in all Dbt-iP3F and Dbt-MYCN-iP3F lines (Fig. 2B and S2A). In the second set, exemplified by *FGF8*, the target gene is not stimulated by P3F in Dbt-iP3F-Pa lines but is stimulated by P3F in Dbt-iP3F-TD and in Dbt-MYCN-iP3F-Pa and TD lines (Fig. 2C and S2B).

We previously demonstrated that FGF8 overexpression is oncogenic in a subset of recurrent tumors from the myoblast system and that FGF8 promotes tumor formation in Dbt-MYCN lines in the absence of P3F [22]. We further investigated if FGF8 is sufficient to transform Dbt-iP3F-Pa cells in the absence of added MYC and/or MYCN. For this experiment, we transfected a constitutive *FGF8* expression construct or empty vector [22] into a Dbt-iP3F-Pa line and confirmed high FGF8 protein expression (Fig. 2D). In a focus formation assay of oncogenic transformation, our results indicate that FGF8 expression is sufficient to induce transformation, even in the absence of high-level P3F (Fig. 2E). Next, we addressed whether FGF8 is required for oncogenicity in Dbt-iP3F-TD lines. Using a CRISPR/Cas9 approach to knockdown *FGF8* in Dbt-iP3F-TD lines, we confirmed that the treated cells showed low or no FGF8 expression (Fig. 2F). Focus formation assays then indicated that *FGF8* knockdown in Dbt-iP3F-TD lines impaired transforming activity in the presence of P3F (Fig. 2G), thus indicating that FGF8 is necessary for oncogenicity of this Dbt-iP3F-TD line.

### MYCN is required for P3F to bind to the *FGF8* locus

To further characterize the biological role of MYCN in P3F-induced transcriptional activation of *FGF8*, we performed chromatin immunoprecipitation sequencing (ChIP-seq) experiments in the P3F-inducible myoblast lines with and without MYCN at various time points following doxycycline addition (0, 8 and 24 hours). We focused on two known P3F targets: *FGFR4*, which does not require MYCN for P3F-induced upregulation (Fig. 2B and S2A) and *FGF8*, which requires MYCN for upregulation (Fig. 2C and S2B). Our ChIP-seq analysis revealed strong enrichment of P3F binding in an *FGF8* intron region (5 kb downstream of its promoter) in MYCN-expressing Dbt-MYCN-iP3F-Pa cells (∼120-fold increase in ChIP enrichment upon P3F induction [p=0.00106]), and weaker P3F binding in MYCN-deficient Dbt-iP3F-Pa cells (30-fold increase in ChIP enrichment upon P3F induction [p=0.0734]) (Fig. 3A and 3C). This finding contrasts to the P3F occupancy in the two regulatory regions of *FGFR4*; at the more proximal site, there is 80-fold and 130-fold increases in ChIP enrichment in Dbt-iP3F and Dbt-MYCN-iP3F cells, respectively, upon P3F induction (Fig. 3B and 3C). These results, combined with our gene expression analysis, support the premise that MYCN is required for P3F to bind at a level that can transcriptionally activate high *FGF8* expression.

**Figure 3.**
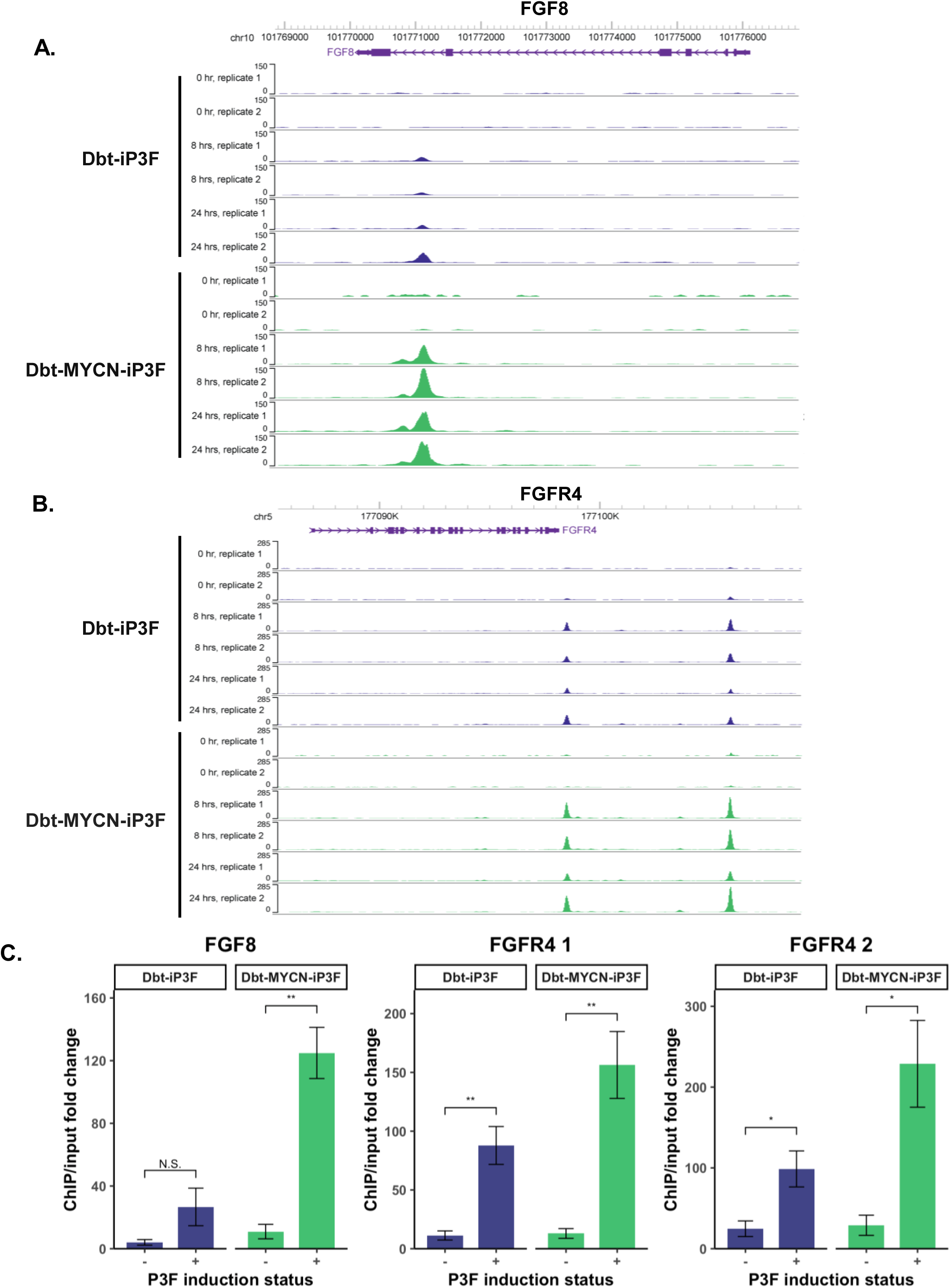
Chromatin immunoprecipitation (ChIP) analysis of P3F binding to FGF8 and FGFR4 loci. **A, B.** P3F ChIP-seq signal at *FGF8* (A) and *FGFR4* (B) in Dbt-iP3F-Pa and Dbt-MYCN-iP3F-Pa cells after doxycycline addition (500 ng/ml) for 0, 8, or 24 hours. **C.** ChIP fold-change compared to input for P3F peaks at *FGF8* and *FGFR4* loci in parts A and B. FGFR4 1 and FGFR4 2 are the more proximal and more distal peaks, respectively, relative to the FGFR4 gene body. Each measurement is expressed as the mean ± SE, n= 4 for +P3F induction and n=2 for -P3F induction. Welch’s t-test was used to compare conditions. Significance levels are described in Fig. 1.

### Dependence of myoblast lines on MYC and MYCN for growth and oncogenicity

We next investigated whether MYC and MYCN contribute to growth and oncogenicity of the engineered myoblast lines. For these experiments, we used a CRISPR/Cas9 (CR) loss-of-function strategy to knockdown *MYC* or *MYCN* in the Dbt-iP3F-TD lines and Dbt-MYCN-iP3F-Pa and TD lines. Western blot analysis after Cas9/gRNA transduction showed 70-90% decrease in the MYC or MYCN protein level (48 hours after transduction without antibiotic selection) (Fig. 4A and 4D).

**Figure 4.**
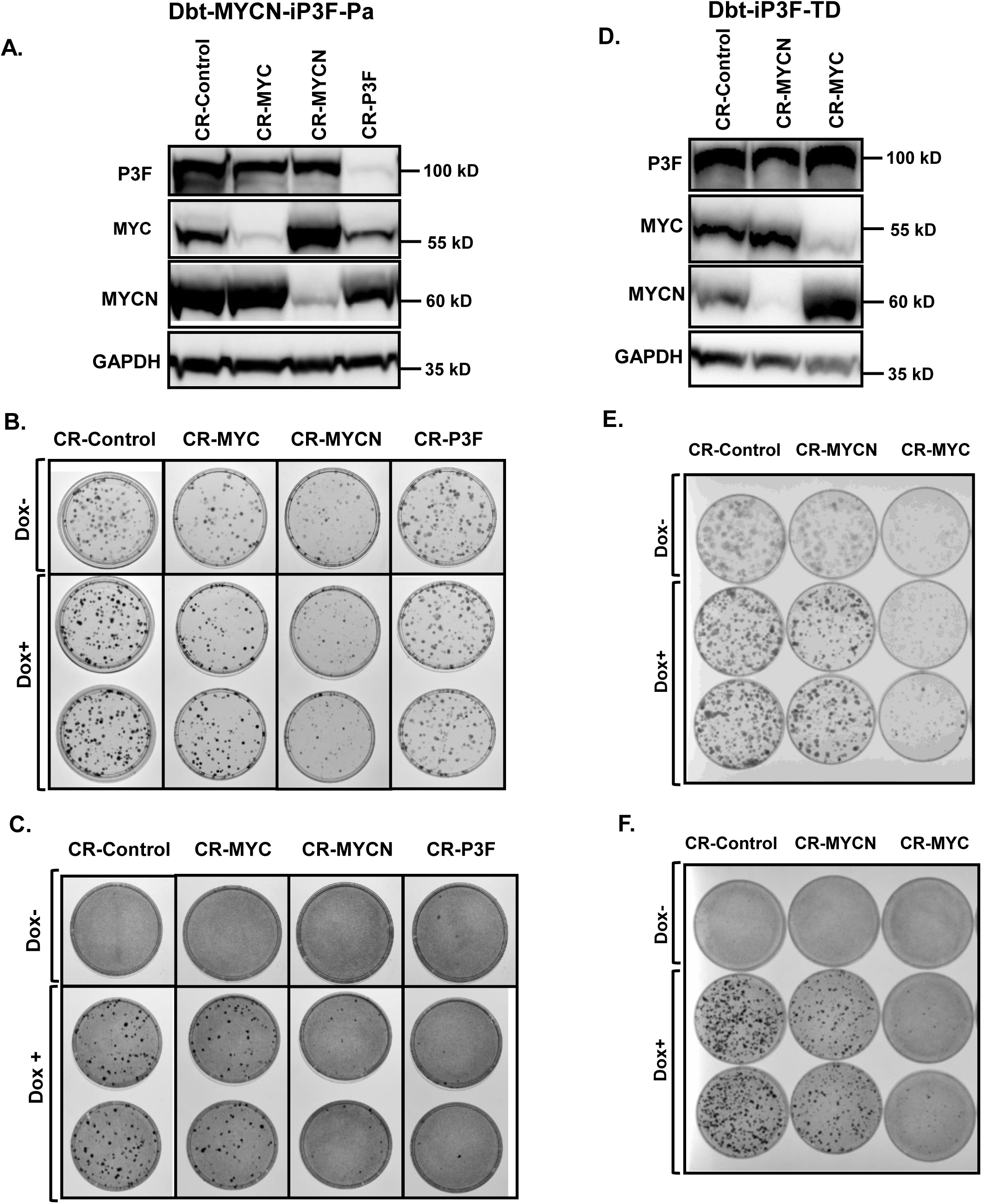
*MYC* and *MYCN* knockdown in engineered myoblast lines. **A, B,C.** Western blot analysis of P3F, MYC and MYCN protein expression (A), clonogenic assay (B) and focus formation assay (C) in Dbt-MYCN-iP3F-Pa line after CRISPR/Cas9 (CR) knockdown of *MYC*, *MYCN* or *P3F*. **D, E, F.** Western blot analysis of P3F, MYC and MYCN protein expression (D), clonogenic assay (E) and focus formation assay (F) in Dbt-iP3F-TD line after CRISPR/Cas9 knockdown of *MYC* or *MYCN*. Clonogenic and focus formation assays were conducted in the presence (+) and absence (-) of 500 ng/ml doxycycline (Dox). See Figure S3 for quantitative results of clonogenic and focus formation assays.

In a Dbt-MYCN-iP3F Pa line, knockdown of *MYCN* resulted in substantial loss of clonogenic growth and focus formation while knockdown of *MYC* resulted in a lesser effect (Fig. 4B-C and S3D-E), indicating that MYCN is the main Myc family member required for growth and oncogenicity in these cells. Of note, P3F knockdown in these cells eliminated focus formation, but still permitted clonogenic growth. It should also be noted that *MYCN* knockdown resulted in upregulation of MYC (Fig. 4A), but this increased MYC level did not rescue the growth and oncogenicity of these cells (Fig. 4B-C). A similar experiment with a Dbt-MYCN-iP3F-TD line also revealed a greater dependence of clonogenic growth and focus formation on MYCN compared to MYC (Fig. S3A-C, S3H-I).

Similar knockdown studies were also performed in a Dbt-iP3F-TD line. In this line, *MYC* knockdown significantly decreased the ability of these cells for clonogenic growth and focus formation whereas *MYCN* knockdown only partially reduced the growth and transforming ability of these cells (Fig. 4D-F and S3F-G). Although *MYC* knockdown resulted in increased MYCN expression (Fig. 4D), this increased MYCN level did not rescue the growth and oncogenicity of these cells (Fig. 4E-F). These results indicate that MYC is the Myc family member primarily responsible for growth and oncogenicity in these cells.

### Dependence of FP-RMS lines on MYC and MYCN for growth and oncogenicity

To elucidate the role of MYC and MYCN in the growth and oncogenic transformation of FP-RMS cell lines, we also used a CRISPR/Cas9 system to knockdown specific genes in these cell lines. In addition to individual *MYC* or *MYCN* knockdown, we utilized a combined gRNA construct to simultaneously knockdown *MYC* and *MYCN* expression in these FP-RMS lines. FP-RMS cell lines were selected with puromycin for 48 hours following 24 hours of transduction with CRISPR/Cas9 constructs with puromycin resistant gene targeting P3F, MYC and/or MYCN. As a positive control in these studies, knockdown of the specific *P3F* or *P7F* fusion gene in each cell line diminished the cell growth and oncogenic properties in all three subsets of FP-RMS cell lines (Fig. 5, S4, S5 and S6). Of note, fusion gene knockdown in these RMS lines resulted in decreased MYCN and variable changes in MYC expression in addition to decreasing fusion protein expression.

**Figure 5.**
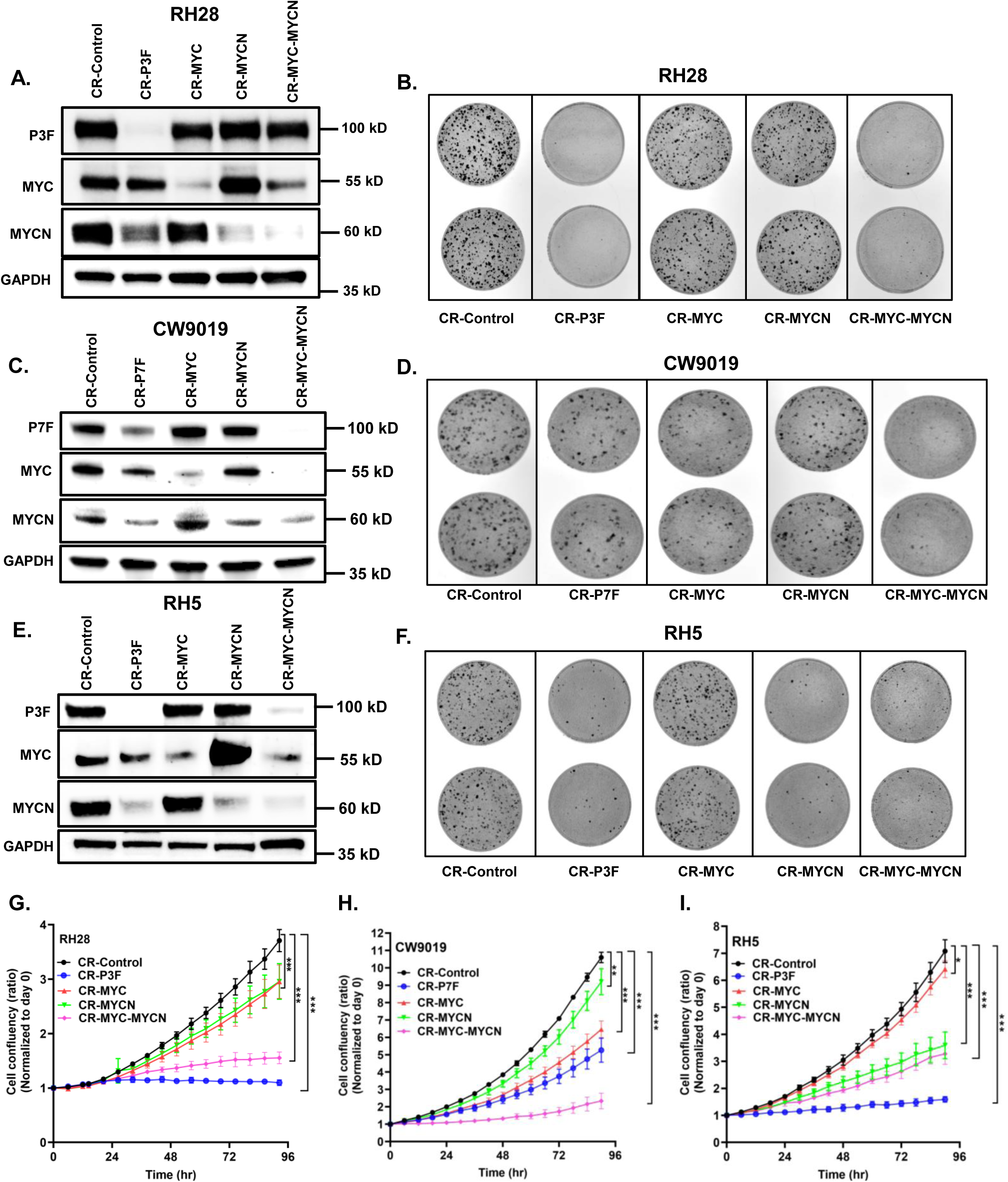
*MYC* and *MYCN* knockdown in FP-RMS lines. **A, B.** Western blot analysis of P3F, MYC and MYCN expression (A) and focus formation assay (B) in RH28 cells after CRISPR/Cas9 (CR) knockdown of *P3F*, *MYC* and/or *MYCN*. **C, D.** Western blot analysis of P7F, MYC and MYCN expression (C) and focus formation assay (D) in CW9019 cells after CRISPR/Cas9 knockdown of *P7F*, *MYC* and/or *MYCN*. **E, F.** Western blot analysis of P3F, MYC and MYCN expression (E) and focus formation assay (F) in RH5 cells after CRISPR/Cas9 knockdown of *P3F*, *MYC* and/or *MYCN*. **G, H, I.** IncuCyte assay of population growth in Rh28 (G), CW9019 (H) and RH5 (I) cells after CRISPR/Cas9 knockdown of *P3F/P7F*, *MYC* and/or *MYCN*. Significance levels are described in Fig. 1. See Figures S4, S5 and S6 for quantitative results of focus formation assays in RH28, CW9019 and RH5, respectively.

In the *P3F*-positive RH28 line, which is an example of a MYC/MYCN co-dominant line, knockdown of either *MYC* or *MYCN* resulted in partial loss of oncogenic transformation (Fig. 5A-B and S4C). Furthermore, there was also a partial inhibition of population cell growth and clonogenic growth with *MYC* or *MYCN* knockdown in RH28 cells when compared with CRISPR-control cells (Fig. 5G and S4A-B). In contrast, double knockdown of *MYC* and *MYCN* in RH28 cells resulted in a near complete loss of cell growth, clonogenicity and oncogenic transformation, similar to the results of the *P3F* knockdown (Fig. 5A-B, 5G and S4). Comparable results for the role of MYC and/or MYCN were seen in RH30, which is another MYC/MYCN co-dominant line (data not shown).

In the *P7F*-positive CW9019 line, which is an example of a MYC-dominant line, *MYC* knockdown attenuated MYC protein expression and significantly diminished oncogenic transformation (Fig. 5C-D and S5C). A reciprocal increase in MYCN expression in response to *MYC* knockdown did not rescue this phenotypic effect. Similarly, population and clonogenic growth were also markedly reduced with *MYC* knockdown in CW9019 cells (Fig. 5H and S5A-B). In contrast, *MYCN* knockdown only showed a small inhibitory effect on these phenotypic parameters in CW9019 cells. Furthermore, double knockdown of *MYC* and *MYCN* in CW9019 cells further impaired oncogenic transformation, population growth and clonogenicity when compared to *MYC* knockdown in these cells (Fig. 5C-D, 5H and S5). These double knockdown results suggest that MYCN partially contributes to these oncogenic properties in CW9019 cells; part of this effect may be related to the complete loss of P7F expression resulting from combined *MYC/MYCN* knockdown in these cells.

In the *P3F*-positive RH5 line, which is an example of a MYCN-dominant line, *MYCN* knockdown significantly diminished MYCN expression accompanied by a marked upregulation of MYC expression (Fig. 5E). In addition to decreasing the protein levels of both MYC and MYCN, double *MYC/MYCN* knockdown also substantially reduced P3F expression. Whereas *MYC* knockdown had little or no effect on focus formation, population growth and clonogenicity of RH5 cells, *MYCN* knockdown produced a strong inhibition of these phenotypic parameters (Fig. 5F, 5I and S6A-C). Unlike the findings in RH28 and CW9019, the phenotype of the double *MYC/MYCN* knockdown was comparable to that of the *MYCN* knockdown in RH5 cells; this finding suggests that the amplified *MYCN* gene provides all the Myc family protein function and MYC does not play much role in regulating growth and oncogenicity in RH5 cells. Even the MYC upregulation in response to *MYCN* knockdown does not rescue the growth and oncogenicity of these cells.

### Role of Myc family proteins in maintaining P3F target expression

We established that MYCN is required for P3F to optimally upregulate *FGF8* in the engineered myoblasts whereas this Myc protein is not needed for P3F upregulation of most other target genes, such as *FGFR4*. We next investigated whether MYCN is required for continual expression of these P3F target genes in Dbt-MYCN-iP3F lines. RNA-Seq analysis of three Dbt-MYCN-iP3F-Pa lines revealed that expression of *FGF8* and *FGFR4* remains intact following *MYCN* or *MYC* knockdown (Fig. 6A-B) despite the fact that *MYCN* knockdown caused major changes in clonogenic growth and transformation (Fig. 4 and S3). In contrast, both *FGF8* and *FGFR4* expression substantially decrease upon P3F knockdown in the presence of doxycycline. From these results we infer that MYCN is initially required for transcriptional activation of *FGF8*, but once it is activated, high MYCN expression is no longer required to maintain P3F-induced expression.

**Figure 6.**
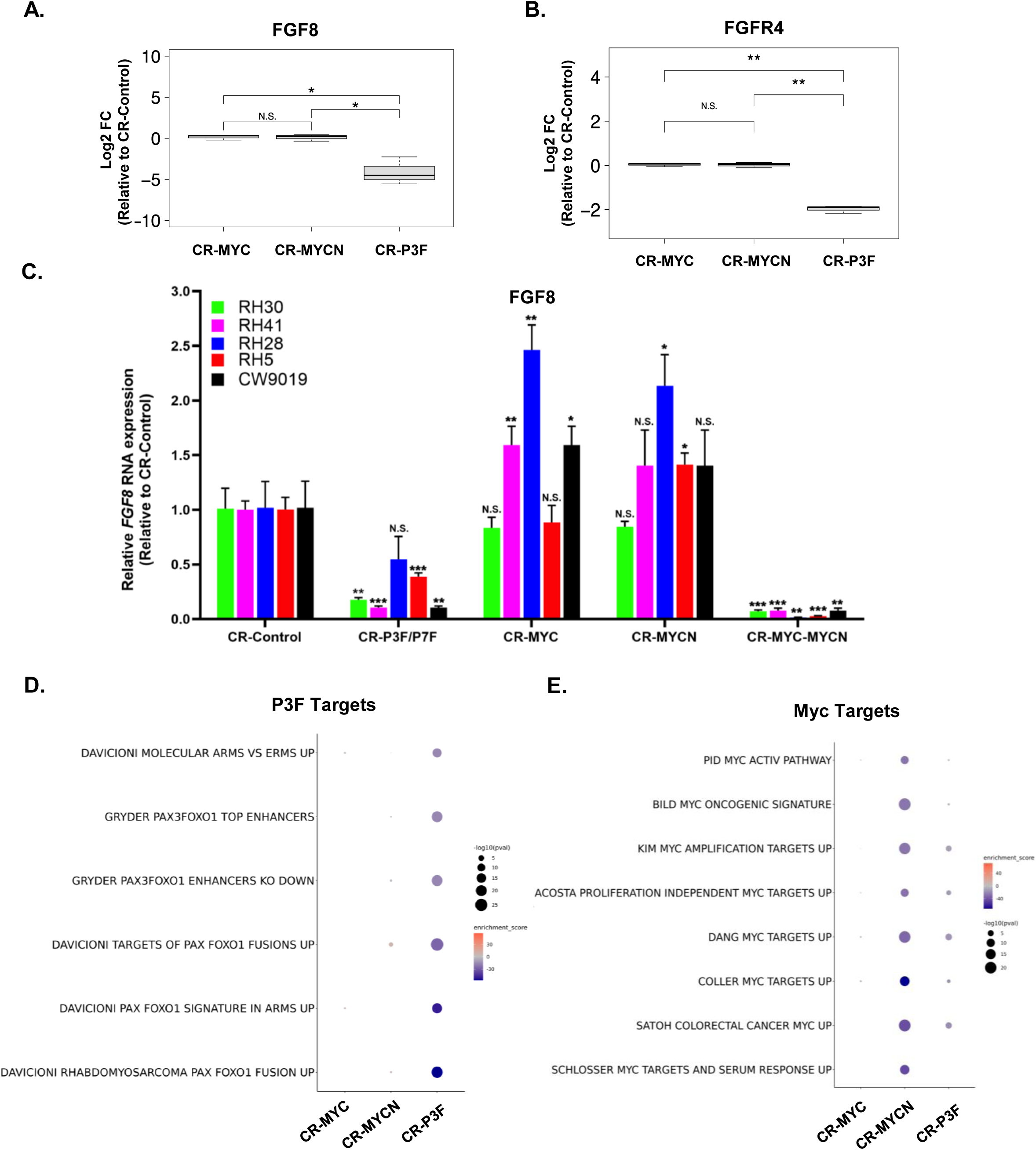
P3F and Myc target gene expression following knockdown of *MYCN* and/or *MYC*. **A, B.** RNA-seq analysis of *FGF8* (A) and *FGFR4* (B) RNA expression in Dbt-MYCN-iP3F myoblasts following CRISPR/Cas9 (CR) knockdown of *P3F*, *MYCN* or *MYC*. Each measurement is expressed as mean ± SE of 3 biological replicates (subclones). **C.** Quantitative RT-PCR (qRT-PCR) analysis of *FGF8* RNA expression in FP-RMS cell lines following CRISPR/Cas9 knockdown of *P3F/P7F*, *MYC* and/or *MYCN*. The *FGF8* RNA levels were normalized with respect to RNA expression in the sample with empty vector (CR-control). Significance levels are described in Fig. 1. **D, E.** Gene set enrichment analysis of P3F (D) and Myc (E) target enrichment in Dbt-MYCN-iP3F-Pa myoblasts following *MYC, MYCN* and *P3F* knockdown. The P-Value and enrichment scores are indicated by the scales shown to the right of the bubble plots.

We further explored whether Myc family proteins regulate *FGF8* RNA expression in FP-RMS cell lines (Fig. 6C). As a positive control, knockdown of *P3F* or *P7F* reduced *FGF8* mRNA levels in all three subsets of FP-RMS cell lines. In contrast, knockdown of either *MYC* or *MYCN* did not reduce the *FGF8* transcript levels despite the fact that *MYCN* knockdown in RH5 and *MYC* knockdown in CW9019 result in decreased growth and transformation. In fact, the *FGF8* mRNA level actually increased in several lines in response to *MYC* or *MYCN* knockdown. Of note, *MYC/MYCN* double knockdown significantly reduced the *FGF8* RNA level in all subsets of FP-RMS cell lines; this effect might be partly attributable to the reduced P3F or P7F protein levels in CW9019 and RH5 (but not RH28) following double *MYC/MYCN* knockdown (Fig. 5A, 5C, 5E). These results indicate that the dominant Myc protein is needed to maintain growth and transformation but is not needed to maintain transcriptional activation of *FGF8*. Furthermore, the increases in *FGF8* mRNA resulting from *MYC* or *MYCN* knockdown did not prevent the decrease in growth and oncogenic properties in these FP-RMS cell lines.

Finally, we compared global downstream effects of *P3F* and *MYCN* knockdown in the Dbt-MYCN-iP3F-Pa myoblast system by performing RNA sequencing and gene set enrichment analysis. We used the C2 curated gene set pathway database from MSigDB (v6.2) [32] to compare P3F- and Myc protein-mediated expression pathways across the individual CRISPR/Cas9 knockdowns of *MYC*, *MYCN* and *P3F* in the Dbt-MYCN-iP3F-Pa line. For P3F-related targets (Fig. 6D and Supp. Table 2), *P3F* knockdown significantly depleted these signatures (FDR<0.0001) while both *MYC* and *MYCN* knockdowns had no effect (FDR=1). For the Myc protein-related targets (Fig. 6E and Supp. Table 3), *MYCN* knockdown and to a lesser degree *P3F* knockdown depleted these signatures. In contrast, these Myc protein-related pathways were not affected by *MYC* knockdown, presumably due to the high level MYCN expression in these cells (FDR=1). These results indicate that Myc family proteins were not necessary for maintenance of expression of up-regulated P3F targets in the Dbt-MYCN-iP3F-Pa cells.

## Discussion

In this manuscript, we explored the expression and function of Myc family proteins in the pathogenesis of FP-RMS. In our study, we noted evidence of an inverse relationship between MYCN and MYC expression in some RMS lines and MYC/MYCN co-expression in other lines. For example, MYCN and MYC levels are low in the MYC-dominant and MYCN-dominant FP-RMS cell lines, respectively, and knockdown of high level MYCN or MYC expression resulted in upregulation of MYC or MYCN levels, respectively. Similarly, expression of exogenous MYCN in Dbt myoblasts resulted in decreased endogenous MYC expression. Differential expression of Myc and Mycn has been previously observed during developmental studies [33, 34] that found Mycn is predominantly expressed during early embryogenesis and is downregulated during adulthood [35] whereas Myc functions in tissues with proliferative capacity during adulthood [33]. An inverse correlation of MYC and MYCN expression at both the RNA and protein levels has been identified in cancers such as neuroblastoma [36, 37] and small cell lung carcinoma [38]. These findings indicate that MYC and MYCN expression is often mutually exclusive and suggest inverse feedback mechanisms in some cell types. However, our finding of FP-RMS cell lines in which MYC and MYCN are co-expressed at high levels suggests that there are also mechanisms in certain cell types that permit co-expression of these two genes.

The *MYC* and *MYCN* genes are often highly expressed and deregulated in different cancer types leading to aberrant activation of Myc signaling pathways, which favor growth and oncogenicity [6]. In neuroblastomas, there is a subset of tumors with *MYCN* gene amplification, which results in high MYCN expression at the RNA and protein levels [36, 37, 39]. We and other previously found *MYCN* gene amplification associated with high *MYCN* expression in a subset of FP-RMS tumors [8]. In the current study, we found *MYCN* gene amplification and high MYCN expression in the MYCN-dominant FP-RMS cell line RH5. Therefore, gene amplification is one mechanism that drives high MYCN expression in pediatric tumors, such as FP-RMS.

Various mechanisms have also been identified for high MYC expression in tumors [40]. *MYC* gene amplification has been detected in several cancer types, including ovarian, esophageal, uterine and breast carcinoma [6]. Moreover, in neuroblastomas without *MYCN* gene amplification, there is a subset with focal amplification of distal enhancers downstream of the *MYC* gene and a second subset with chromosomal translocations that move enhancers from other chromosomes into the vicinity of the *MYC* gene [37]. To date, there is no evidence for these genomic mechanisms in FP-RMS but further analysis may be useful if directed specifically to MYC-dominant FP-RMS tumors.

In our myoblast model system, we found that MYCN collaborates with P3F to upregulate a subset of P3F target genes, including *FGF8*, and thereby induces oncogenic effects. There is published evidence of multiple mechanisms by which Myc family proteins can influence gene expression. First, MYC is a transcription factor that can bind specifically to E-boxes motifs (CACGTG) [41–43] and transactivate target gene expression. In addition to these high affinity sites, Myc can be recruited to genes with low affinity Myc binding sites to “amplify” the existing level of gene expression [44 , 45]. Aside from these DNA binding site-related effects, MYCN can form complexes with transcription factors that bind to other loci and modify chromatin structure to modulate gene expression [46]. A recent study also described the ability of MYCN to interact with the TFIIIC complex and change RNA polymerase II dynamics by changing chromatin architecture [47].

Our analysis of the role of Myc proteins in FP-RMS must also account for the finding that knockdown of dominant Myc proteins inactivates oncogenic effects without decreasing P3F targets such as *FGF8*. One possible explanation is that Myc proteins may initially cause the above-described changes in local chromatin structure at P3F target loci such as *FGF8*, thereby allowing P3F binding and upregulated target gene expression. Once this modification occurs, the chromatin structure may remain open even if the causative Myc protein is removed. Alternatively, the reciprocal upregulation of MYC or MYCN when dominant MYCN or MYC, respectively, is knocked down may be sufficient to maintain the Myc regulatory effect on the *FGF8* locus. This hypothesis is supported by the finding that *FGF8* is silenced when both Myc proteins are knocked down.

Even though Myc protein knockdown does not reverse P3F target gene expression, this knockdown still reverses oncogenic effects in the myoblast system and FP-RMS lines. We postulate that the effect of knockdown of Myc family proteins on growth and oncogenic properties of these cells is mediated through Myc protein pathways unrelated to P3F pathways in these cells. The Myc targets downregulated by MYCN knockout include ribosomal biogenesis and cell cycle regulatory genes and have been identified in other cancer types, including colorectal and small cell lung carcinomas [48–53].

Our studies of co-dominant FP-RMS lines expressing both MYC and MYCN provide evidence that these Myc proteins have overlapping functions. The premise that MYC and MYCN exert redundant functions is supported by developmental studies in which germline knockdown of *Myc* is rescued by transgenic insertion of *Mycn* [54] and conditional deletion of *Mycn* is rescued by *Myc* [55]. The shared functions of these Myc proteins during development have an impact on growth promotion. This common growth-promoting function also extends to many types of cancers with overexpression of Myc proteins [56]. Increased expression of these Myc proteins contribute to oncogenicity in cell line studies [57–60], and depletion of Myc proteins reduces growth in numerous cancer types.

In conclusion, these studies demonstrate that Myc family proteins play an important role in FP-RMS, both by collaborating with the P3F and P7F fusion proteins as well as by activating independent growth-related pathways. There are FP-RMS subsets in which MYCN or MYC is the dominant Myc protein whereas both Myc proteins are co-dominant in other FP-RMS subsets. In addition to providing keys for understanding oncogenic mechanisms in FP-RMS, the Myc family proteins therefore represent an important set of potential therapeutic targets.

## Acknowledgments

This research was supported by the Intramural Research Program of the National Institutes of Health (NIH) and the Joanna McAfee Childhood Cancer Foundation. The contributions of the NIH authors were made as part of their official duties as NIH federal employees, are in compliance with agency policy requirements, and are considered Works of the United States Government. However, the findings and conclusions presented in this paper are those of the authors and do not necessarily reflect the views of the NIH or the U.S. Department of Health and Human Services.

## Notes

### Competing Interest Statement

The authors have declared no competing interest.

## References

1 Ognjanovic S, Linabery AM, Charbonneau B, Ross JA. Trends in childhood rhabdomyosarcoma incidence and survival in the United States, 1975-2005. Cancer 2009; 115: 4218–4226.

2 Skapek SX, Ferrari A, Gupta AA, Lupo PJ, Butler E, Shipley J, et al. Rhabdomyosarcoma. Nat Rev Dis Primers 2019; 5: 1.

3 Galili N, Davis RJ, Fredericks WJ, Mukhopadhyay S, Rauscher FJ, Emanuel BS et al. Fusion of a fork head domain gene to PAX3 in the solid tumour alveolar rhabdomyosarcoma. Nature Genetics 1993; 5: 230–235.

4 Barr FG. Gene fusions involving PAX and FOX family members in alveolar rhabdomyosarcoma. Oncogene 2001; 20: 5736–5746.

5 Whitfield JR, Soucek L. MYC in cancer: from undruggable target to clinical trials. Nat Rev Drug Discov 2025; 24: 445–457.

6 Schaub FX, Dhankani V, Berger AC, Trivedi M, Richardson AB, Shaw R et al. Pan-cancer Alterations of the MYC Oncogene and Its Proximal Network across the Cancer Genome Atlas. Cell Syst 2018; 6: 282–300.e282.

7 Barr FG, Duan F, Smith LM, Gustafson D, Pitts M, Hammond S, Gastier-Foster JM. Genomic and clinical analyses of 2p24 and 12q13-q14 amplification in alveolar rhabdomyosarcoma: a report from the Children’s Oncology Group. Genes Chromosomes Cancer 2009; 48: 661–672.

8 Hachitanda Y, Toyoshima S, Akazawa K, Tsuneyoshi M. N-myc gene amplification in rhabdomyosarcoma detected by fluorescence in situ hybridization: its correlation with histologic features. Mod Pathol 1998; 11: 1222–1227.

9 Mercado GE, Xia SJ, Zhang C, Ahn EH, Gustafson DM, Laé M et al. Identification of PAX3-FKHR-regulated genes differentially expressed between alveolar and embryonal rhabdomyosarcoma: focus on MYCN as a biologically relevant target. Genes Chromosomes Cancer 2008; 47: 510–520.

10 Williamson D, Lu YJ, Gordon T, Sciot R, Kelsey A, Fisher C et al. Relationship between MYCN copy number and expression in rhabdomyosarcomas and correlation with adverse prognosis in the alveolar subtype. J Clin Oncol 2005; 23: 880–888.

11 Cao L, Yu Y, Bilke S, Walker RL, Mayeenuddin LH, Azorsa DO et al. Genome-wide identification of PAX3-FKHR binding sites in rhabdomyosarcoma reveals candidate target genes important for development and cancer. Cancer Res 2010; 70: 6497–6508.

12 Xia SJ, Holder DD, Pawel BR, Zhang C, Barr FG. High expression of the PAX3-FKHR oncoprotein is required to promote tumorigenesis of human myoblasts. Am J Pathol 2009; 175: 2600–2608.

13 Pandey PR, Chatterjee B, Olanich ME, Khan J, Miettinen MM, Hewitt SM, Barr FG. PAX3-FOXO1 is essential for tumour initiation and maintenance but not recurrence in a human myoblast model of rhabdomyosarcoma. J Pathol 2017; 241: 626–637.

14 Livak KJ, Schmittgen TD. Analysis of relative gene expression data using real-time quantitative PCR and the 2(-Delta Delta C(T)) Method. Methods 2001; 25: 402–408.

15 Boudjadi S, Kim H, Chatterjee B, Raut PK, Nguyen TH, Pandey PR et al. Involvement of the FGF8/FGF receptor signaling pathway in the maintenance and progression of fusion-positive rhabdomyosarcoma. Mol Cancer Ther 2025 (in press).

16 Martin M. Cutadapt removes adapter sequences from high-throughput sequencing reads. EMBnetjournal 2011; 17: 10--12.

17 Dobin A, Davis CA, Schlesinger F, Drenkow J, Zaleski C, Jha S et al. STAR: ultrafast universal RNA-seq aligner. Bioinformatics 2013; 29: 15–21.

18 Li B, Dewey CN. RSEM: accurate transcript quantification from RNA-Seq data with or without a reference genome. BMC Bioinformatics 2011; 12: 323.

19 Ritchie ME, Phipson B, Wu D, Hu Y, Law CW, Shi W, Smyth GK. limma powers differential expression analyses for RNA-sequencing and microarray studies. Nucleic Acids Res 2015; 43: e47.

20 Law CW, Alhamdoosh M, Su S, Dong X, Tian L, Smyth GK, Ritchie ME. RNA-seq analysis is easy as 1-2-3 with limma, Glimma and edgeR. *F1000Res* 2016; 5.

21 Olanich ME, Sun W, Hewitt SM, Abdullaev Z, Pack SD, Barr FG. CDK4 Amplification Reduces Sensitivity to CDK4/6 Inhibition in Fusion-Positive Rhabdomyosarcoma. Clin Cancer Res 2015; 21: 4947–4959.

22 Boudjadi S, Pandey PR, Chatterjee B, Nguyen TH, Sun W, Barr FG. A Fusion Transcription Factor-Driven Cancer Progresses to a Fusion-Independent Relapse via Constitutive Activation of a Downstream Transcriptional Target. Cancer Res 2021; 81: 2930–2942.

23 Xia SJ, Rajput P, Strzelecki DM, Barr FG. Analysis of genetic events that modulate the oncogenic and growth suppressive activities of the PAX3-FKHR fusion oncoprotein. Lab Invest 2007; 87: 318–325.

24 Schneider CA, Rasband WS, Eliceiri KW. NIH Image to ImageJ: 25 years of image analysis. Nat Methods 2012; 9: 671–675.

25 Sunkel BD, Wang M, LaHaye S, Kelly BJ, Fitch JR, Barr FG et al. Evidence of pioneer factor activity of an oncogenic fusion transcription factor. iScience 2021; 24: 102867.

26 Landt SG, Marinov GK, Kundaje A, Kheradpour P, Pauli F, Batzoglou S et al. ChIP-seq guidelines and practices of the ENCODE and modENCODE consortia. Genome Res 2012; 22: 1813–1831.

27 Li D, Purushotham D, Harrison JK, Hsu S, Zhuo X, Fan C et al. WashU Epigenome Browser update 2022. Nucleic Acids Res 2022; 50: W774–w781.

28 Lawrence M, Huber W, Pagès H, Aboyoun P, Carlson M, Gentleman R et al. Software for computing and annotating genomic ranges. PLoS Comput Biol 2013; 9: e1003118.

29 Wickham H. ggplot2: Elegant Graphics for Data Analysis. Springer-Verlag New York, 2016.

30 Frascella E, Lenzini E, Schafer BW, Brecevic L, Dorigo E, Toffolatti L et al. Concomitant amplification and expression of PAX7-FKHR and MYCN in a human rhabdomyosarcoma cell line carrying a cryptic t(1;13)(p36;q14). Cancer Genet Cytogenet 2000; 121: 139–145.

31 Gryder BE, Yohe ME, Chou HC, Zhang X, Marques J, Wachtel M et al. PAX3-FOXO1 Establishes Myogenic Super Enhancers and Confers BET Bromodomain Vulnerability. Cancer Discov 2017; 7: 884–899.

32 Liberzon A, Subramanian A, Pinchback R, Thorvaldsdóttir H, Tamayo P, Mesirov JP. Molecular signatures database (MSigDB) 3.0. Bioinformatics 2011; 27: 1739–1740.

33 Hurlin PJ. Control of vertebrate development by MYC. Cold Spring Harb Perspect Med 2013; 3: a014332.

34 Jha RK, Kouzine F, Levens D. MYC function and regulation in physiological perspective. Front Cell Dev Biol 2023; 11: 1268275.

35 Ruiz-Pérez MV, Henley AB, Arsenian-Henriksson M. The MYCN Protein in Health and Disease. Genes (Basel) 2017; 8.

36 Upton K, Modi A, Patel K, Kendsersky NM, Conkrite KL, Sussman RT et al. Epigenomic profiling of neuroblastoma cell lines. Sci Data 2020; 7: 116.

37 Zimmerman MW, Liu Y, He S, Durbin AD, Abraham BJ, Easton J et al. MYC Drives a Subset of High-Risk Pediatric Neuroblastomas and Is Activated through Mechanisms Including Enhancer Hijacking and Focal Enhancer Amplification. Cancer Discov 2018; 8: 320–335.

38 Fiorentino FP, Tokgün E, Solé-Sánchez S, Giampaolo S, Tokgün O, Jauset T et al. Growth suppression by MYC inhibition in small cell lung cancer cells with TP53 and RB1 inactivation. Oncotarget 2016; 7: 31014–31028.

39 Seeger RC, Brodeur GM, Sather H, Dalton A, Siegel SE, Wong KY, Hammond D. Association of multiple copies of the N-myc oncogene with rapid progression of neuroblastomas. N Engl J Med 1985; 313: 1111–1116.

40 Dhanasekaran R, Deutzmann A, Mahauad-Fernandez WD, Hansen AS, Gouw AM, Felsher DW. The MYC oncogene - the grand orchestrator of cancer growth and immune evasion. Nat Rev Clin Oncol 2022; 19: 23–36.

41 Blackwell TK, Kretzner L, Blackwood EM, Eisenman RN, Weintraub H. Sequence-specific DNA binding by the c-Myc protein. Science 1990; 250: 1149–1151.

42 Kerkhoff E, Bister K, Klempnauer KH. Sequence-specific DNA binding by Myc proteins. Proc Natl Acad Sci U S A 1991; 88: 4323–4327.

43 Guo J, Li T, Schipper J, Nilson KA, Fordjour FK, Cooper JJ et al. Sequence specificity incompletely defines the genome-wide occupancy of Myc. Genome Biol 2014; 15: 482.

44 Nie Z, Hu G, Wei G, Cui K, Yamane A, Resch W et al. c-Myc is a universal amplifier of expressed genes in lymphocytes and embryonic stem cells. Cell 2012; 151: 68–79.

45 Pellanda P, Dalsass M, Filipuzzi M, Loffreda A, Verrecchia A, Castillo Cano V et al. Integrated requirement of non-specific and sequence-specific DNA binding in Myc-driven transcription. Embo j 2021; 40: e105464.

46 Liu Z, Zhang X, Xu M, Hong JJ, Ciardiello A, Lei H et al. MYCN drives oncogenesis by cooperating with the histone methyltransferase G9a and the WDR5 adaptor to orchestrate global gene transcription. PLoS Biol 2024; 22: e3002240.

47 Vidal R, Leen E, Herold S, Müller M, Fleischhauer D, Schülein-Völk C et al. Association with TFIIIC limits MYCN localisation in hubs of active promoters and chromatin accumulation of non-phosphorylated RNA polymerase II. Elife 2024; 13.

48 Acosta JC, Ferrándiz N, Bretones G, Torrano V, Blanco R, Richard C et al. Myc inhibits p27-induced erythroid differentiation of leukemia cells by repressing erythroid master genes without reversing p27-mediated cell cycle arrest. Mol Cell Biol 2008; 28: 7286–7295.

49 Satoh K, Yachida S, Sugimoto M, Oshima M, Nakagawa T, Akamoto S et al. Global metabolic reprogramming of colorectal cancer occurs at adenoma stage and is induced by MYC. Proc Natl Acad Sci U S A 2017; 114: E7697–e7706.

50 Kim YH, Girard L, Giacomini CP, Wang P, Hernandez-Boussard T, Tibshirani R et al. Combined microarray analysis of small cell lung cancer reveals altered apoptotic balance and distinct expression signatures of MYC family gene amplification. Oncogene 2006; 25: 130–138.

51 Coller HA, Grandori C, Tamayo P, Colbert T, Lander ES, Eisenman RN, Golub TR. Expression analysis with oligonucleotide microarrays reveals that MYC regulates genes involved in growth, cell cycle, signaling, and adhesion. Proc Natl Acad Sci U S A 2000; 97: 3260–3265.

52 Schlosser I, Hölzel M, Hoffmann R, Burtscher H, Kohlhuber F, Schuhmacher M et al. Dissection of transcriptional programmes in response to serum and c-Myc in a human B-cell line. Oncogene 2005; 24: 520–524.

53 Zeller KI, Jegga AG, Aronow BJ, O’Donnell KA, Dang CV. An integrated database of genes responsive to the Myc oncogenic transcription factor: identification of direct genomic targets. Genome Biol 2003; 4: R69.

54 Malynn BA, de Alboran IM, O’Hagan RC, Bronson R, Davidson L, DePinho RA, Alt FW. N-myc can functionally replace c-myc in murine development, cellular growth, and differentiation. Genes Dev 2000; 14: 1390–1399.

55 Muñoz-Martín N, Sierra R, Schimmang T, Villa Del Campo C, Torres M. Myc is dispensable for cardiomyocyte development but rescues Mycn-deficient hearts through functional replacement and cell competition. Development 2019; 146.

56 Lourenco C, Resetca D, Redel C, Lin P, MacDonald AS, Ciaccio R et al. MYC protein interactors in gene transcription and cancer. Nat Rev Cancer 2021; 21: 579–591.

57 Schwab M, Varmus HE, Bishop JM. Human N-myc gene contributes to neoplastic transformation of mammalian cells in culture. Nature 1985; 316: 160–162.

58 Yancopoulos GD, Nisen PD, Tesfaye A, Kohl NE, Goldfarb MP, Alt FW. N-myc can cooperate with ras to transform normal cells in culture. Proc Natl Acad Sci U S A 1985; 82: 5455–5459.

59 Cavalieri F, Goldfarb M. N-myc proto-oncogene expression can induce DNA replication in Balb/c 3T3 fibroblasts. Oncogene 1988; 2: 289–291.

60 Mateyak MK, Obaya AJ, Adachi S, Sedivy JM. Phenotypes of c-Myc-deficient rat fibroblasts isolated by targeted homologous recombination. Cell Growth Differ 1997; 8: 1039–1048.

