## Supplementary Figure S1-S6 for "Role of Myc family proteins in transcriptional regulation of growth and oncogenic transformation in fusion-positive rhabdomyosarcoma"

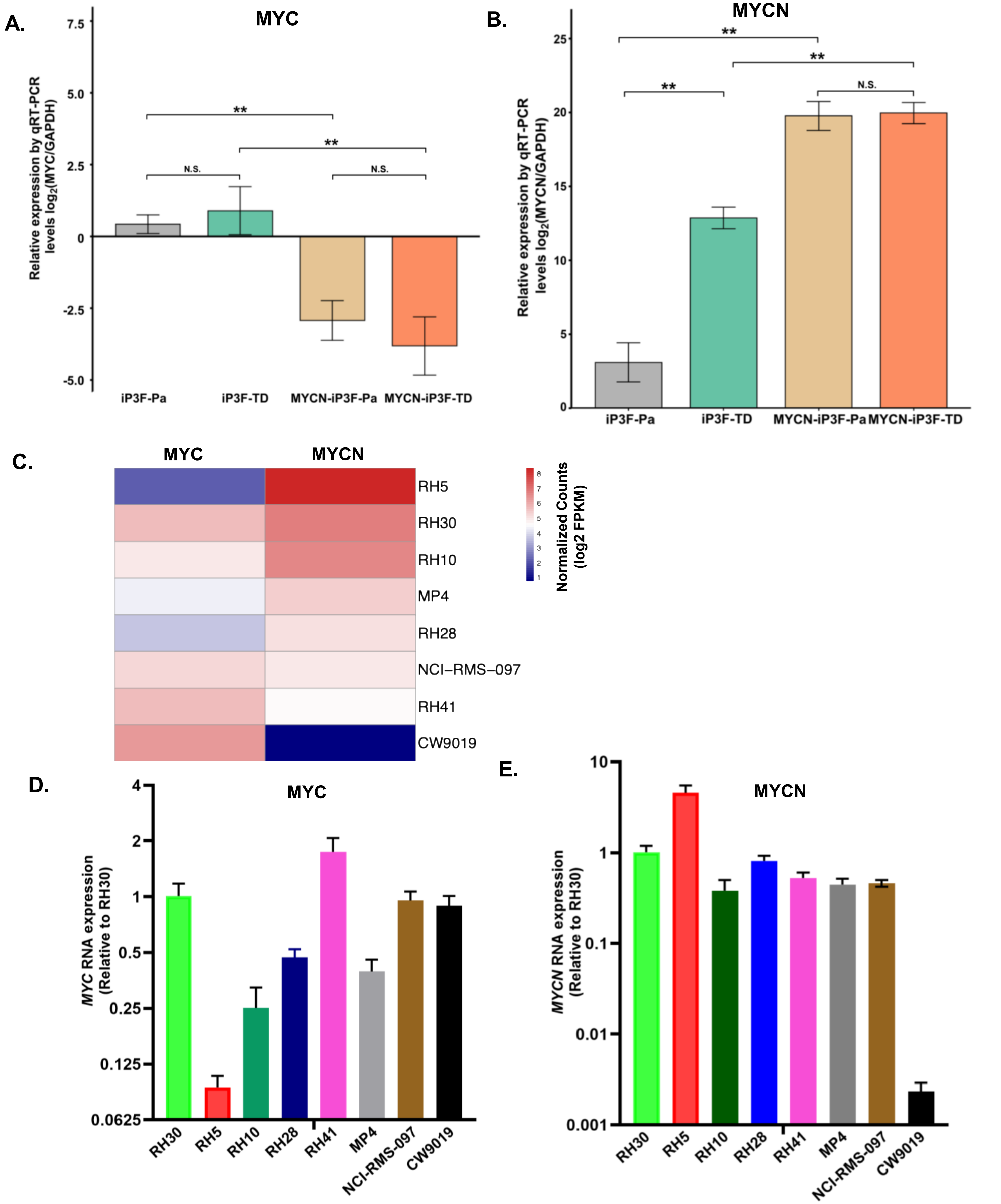


**Supplementary Figure S1. Expression of Myc family members in engineered myoblasts and FP-RMS lines. A, B.** Quantitative RT-PCR (qRT-PCR) analysis of *MYC* (A) and *MYCN* (B) RNA expression in parental (Pa) and tumor-derived (TD) myoblast lines following doxycycline induction (500 ng/ml) of P3F. Significance levels are: *, $P < 0.05$; **, $P < 0.01$; ***, $P < 0.001$; and N.S., not significant. **C.** Heat map of *MYC* and *MYCN* RNA expression in FP-RMS cell lines derived from RNA-Seq data. **D, E.** qRT-PCR analysis of *MYC* (D) and *MYCN* (E) RNA expression in FP-RMS cell lines. The *MYC* and *MYCN* RNA levels were normalized with respect to the expression in RH30 cells.

### Supplementary Figure S2

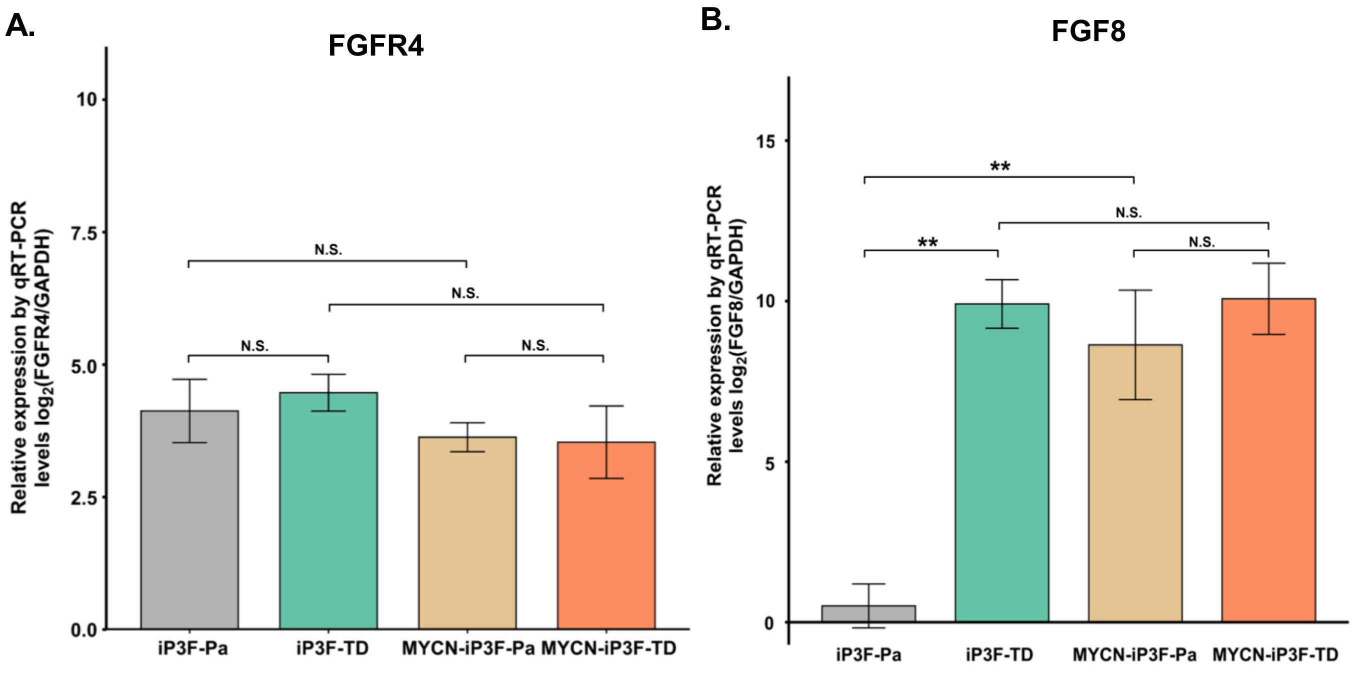


**Supplementary Figure S2. Expression of *FGFR4* and *FGF8* in engineered myoblasts. A, B.** qRT-PCR analysis of *FGFR4* (A) and *FGF8* (B) RNA expression in parental (Pa) and tumor-derived (TD) myoblast lines following doxycycline induction (500 ng/ml) of P3F. Significance levels are described in Fig. S1.

### Supplementary Figure S3

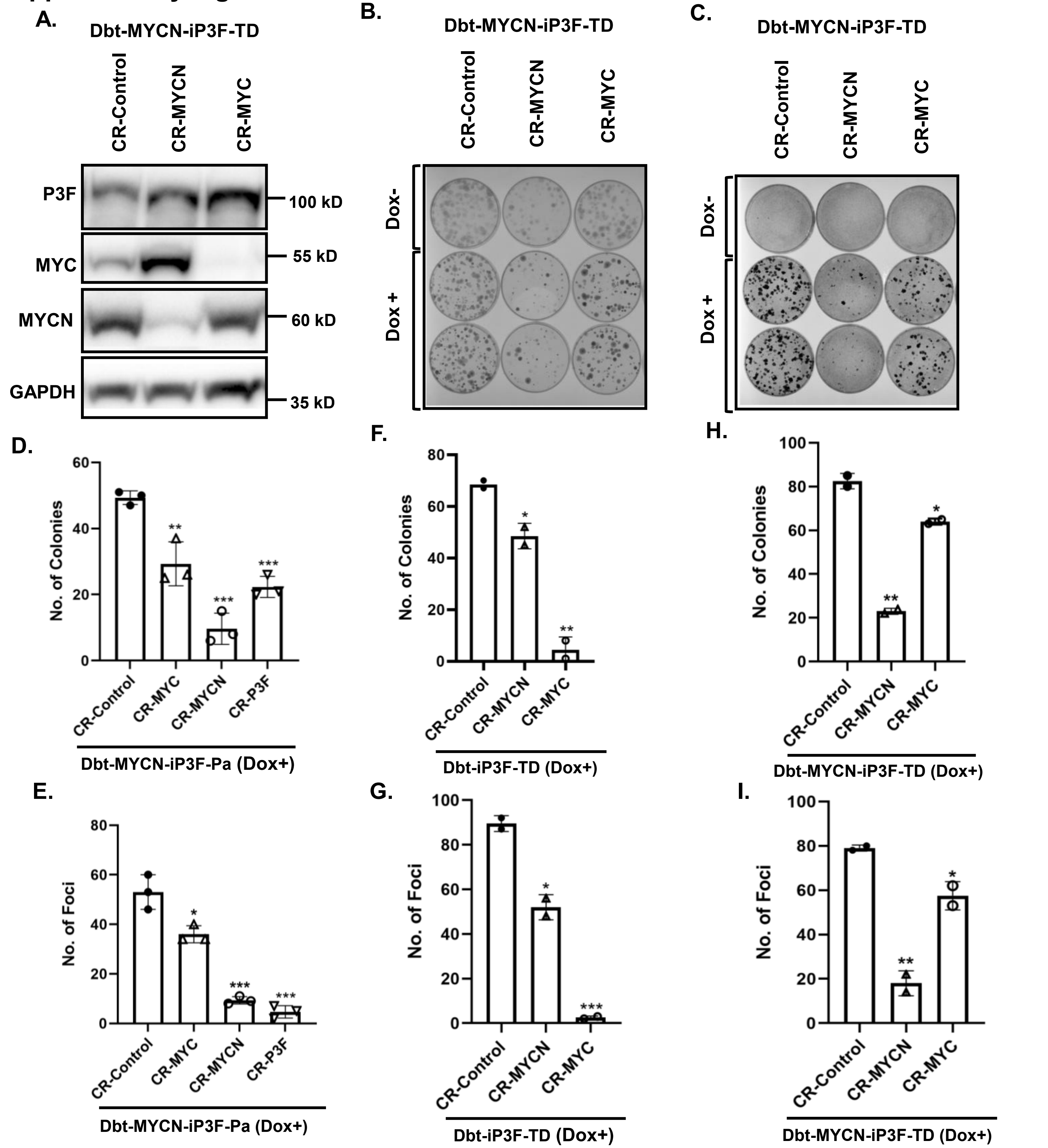


**Supplementary Figure S3. *MYC* and *MYCN* knockdown in engineered myoblast lines. A, B, C.** Western blot analysis of P3F, MYC and MYCN protein expression (A), clonogenic (B), and focus formation (C) assays in a Dbt-MYCN-iP3F-TD line after CRISPR/Cas9 (CR) knockdown of *MYCN* or *MYC*. Assays were performed in the presence (+) or absence (-) of 500 ng/ml doxycycline (Dox). **D, E.** Quantification of clonogenic (D) and focus formation (E) results for Dbt-MYCN-iP3F-Pa cells shown in Fig. 4B and 4C. **F, G.** Quantification of clonogenic (F) and focus formation (G) results for Dbt-iP3F-TD cells shown in Fig. 4E and 4F. Quantitation of clonogenic assay (Fig. S3F) and focus formation (Fig. S3G) was performed with 2 biological replicates for each condition. **H, I.** Quantification of clonogenic (H) and focus formation (I) results for Dbt-MYCN-iP3F-TD cells shown in Fig. S3B and S3C. For B and C, only two dox-induced replicates were available for quantifying clonogenic (H) and focus formation (I) assays; three replicates were used for quantifying the other assay results. Two-sided unpaired student *t* test was used to measure the statistical significance between the CR-Control group and CR-P3F, CR-MYC, or CR-MYCN groups. Significance levels are described in Fig. S1.

### Supplementary Figure S4

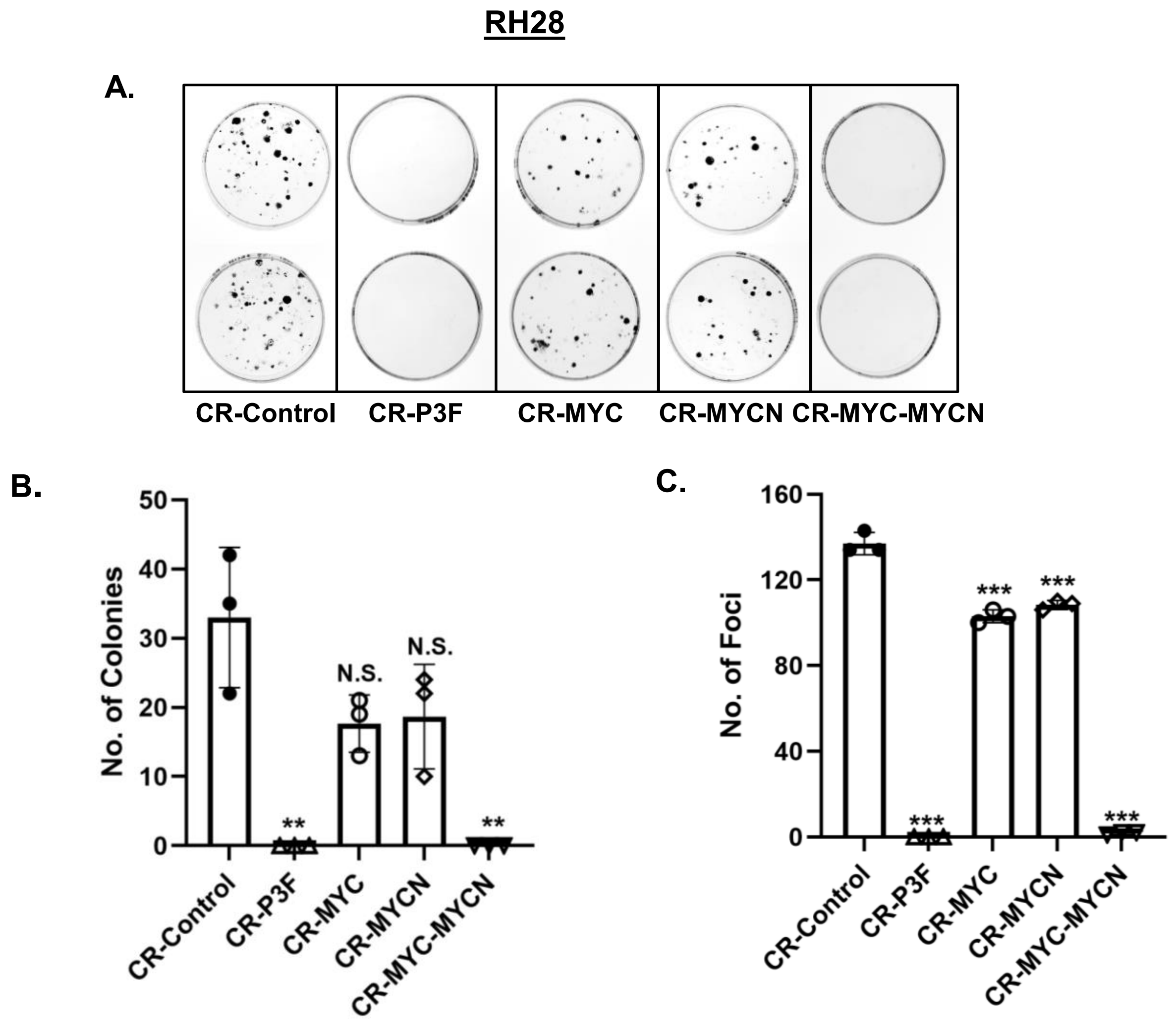


**Supplementary Figure S4:** ***MYC*** **and/or** ***MYCN*** **knockdown in RH28 cells. A.** Clonogenic growth in the RH28 cell line after CRISPR/Cas9 (CR) knockdown of *P3F*, *MYC* and/or *MYCN*. **B, C.** Quantification of clonogenic (B) and focus formation (C) results for RH28 cells shown in Fig. S4A and 5B. A two-sided unpaired student *t* test was used to measure the statistical significance between the CR-Control group and CR-P3F, CR-MYC, CR-MYCN, or CR-MYC-MYCN groups. Significance levels are described in Fig. S1.

### Supplementary Figure S5

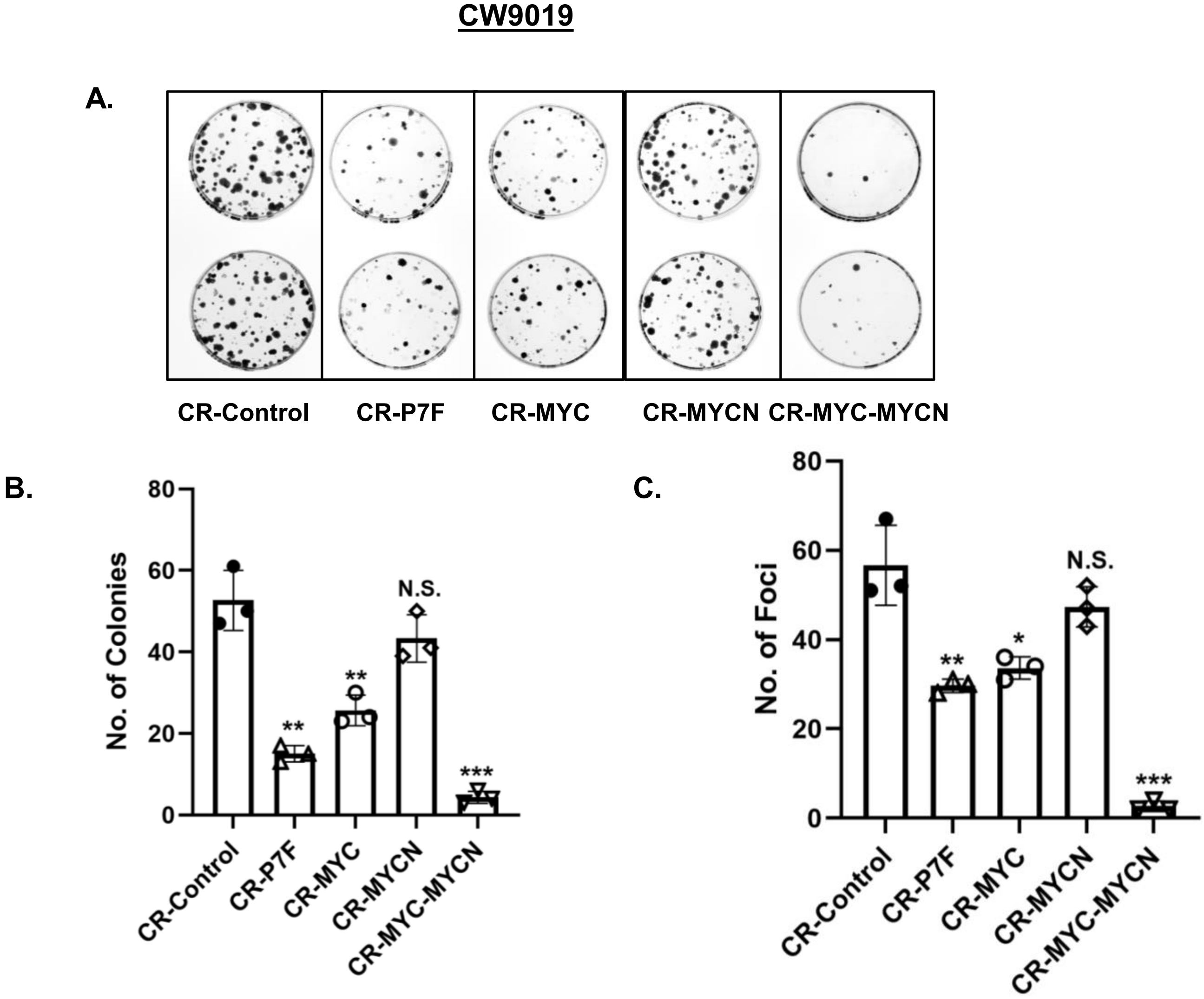


**Supplementary Figure S5: *MYC* and/or *MYCN* knockdown in CW9019 cells. A.** Clonogenic growth in the CW9019 cell line after CRISPR/Cas9 (CR) knockdown of *P7F*, *MYC* and/or *MYCN*. **B, C.** Quantification of clonogenic (B) and focus formation (C) results for CW9019 cells shown in Fig. S5A and 5D. A two-sided unpaired student *t* test was used to measure the statistical significance between the CR-Control group and CR-P7F, CR-MYC, CR-MYCN, or CR-MYC-MYCN groups. Significance levels are described in Fig. S1.

### Supplementary Figure S6

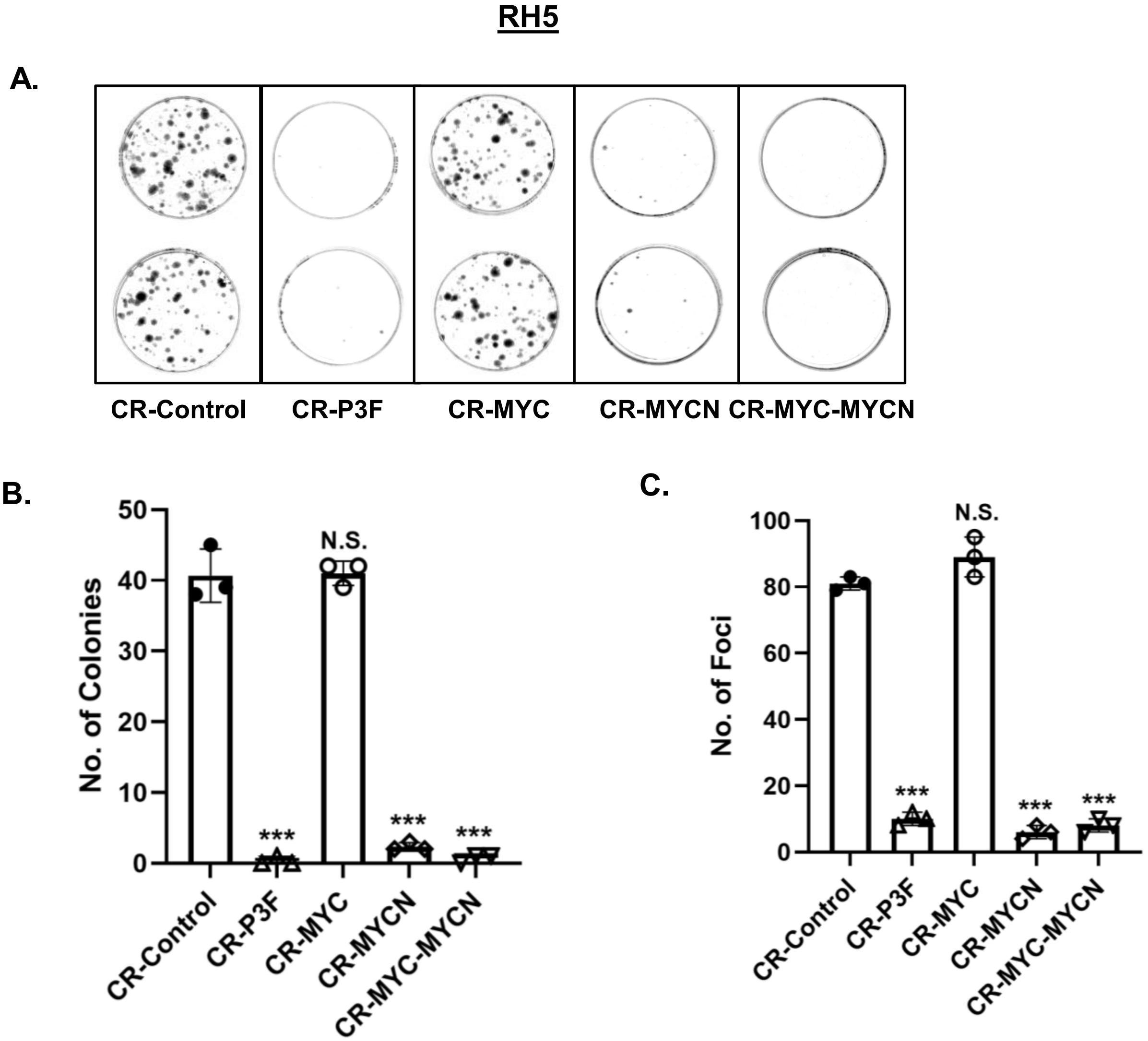


**Supplementary Figure S6: *MYC* and/or *MYCN* knockdown in RH5 cells. A.** Clonogenic growth in the RH5 cell line after CRISPR/Cas9 (CR) knockdown of *P3F*, *MYC* and/or *MYCN*. **B, C.** Quantification of clonogenic (B) and focus formation (C) results for RH5 cells shown in Fig. S6A and 5F. A two-sided unpaired student *t* test was used to measure the statistical significance between the CR-Control group and CR-P3F, CR-MYC, CR-MYCN or CR-MYC-MYCN groups. Significance levels are described in Fig. S1.
