## Supplementary Table 1-3 for "Role of Myc family proteins in transcriptional regulation of growth and oncogenic transformation in fusion-positive rhabdomyosarcoma"

### Supplementary Tables

#### Supplementary Table 1

| Target gene | | gRNA sequence |
|---|---|---|
| **MYC** | CR gRNA-1 | GCCGTATTTCTACTGCGACG |
| | CR gRNA-2 | GTCGAGGTCATAGTTCCTGT |
| **MYCN** | CR gRNA-1 | CACGGCGGGATCCACGCACT |
| | CR gRNA-2 | CAGGCCGAACGCGTCCTCCT |
| **P3F** | CR gRNA-1 | CGCCAGCTGCGCGTGTCCCA |
| **FGF8** | CR gRNA-1 | CAACAAGCGCATCAACGCCA |
| **P7F** | CR gRNA-1 | GGTACCGAGAATGATGCGGC |

**Supplementary Table 1: Guide RNA sequences for different genes**

**Supplementary Table 2**

| Pathway_name | Pathway_Id | Categ | Group | Number_hits | Percent_gene_hits_per_pathway | Enrichment_score | pval | FDR | Net_number_hits | NES |
|---|---|---|---|---|---|---|---|---|---|---|
| DAVICIONI_MOLECULAR_ARMS_VS_ERMS_UP | M2012 | C2 | MYC_KO | 27 | 9.92647059 | 1.39125203 | 0.237113 | 1 | 27 | 1.39125203 |
| | | | MYCN_KO | 29 | 10.6617647 | 0.22120949 | 0.48286989 | 1 | 29 | 0.22120949 |
| | | | P3F_KO | 71 | -26.102941 | -16.300372 | 3.24E-14 | 2.01E-12 | -71 | -16.300372 |
| DAVICIONI_RHABDOMYOSARCOMA_PAX_FOXO1_FUSION_UP | M12362 | C2 | MYC_KO | NA | NA | NA | NA | NA | NA | NA |
| | | | MYCN_KO | 7 | 14.2857143 | 3.85600696 | 0.24737122 | 0.79420558 | 7 | 3.85600696 |
| | | | P3F_KO | 33 | -67.346939 | -57.407842 | 7.92E-22 | 1.50E-19 | -33 | -57.407842 |
| DAVICIONI_TARGETS_OF_PAX_FOXO1_FUSIONS_UP | M4680 | C2 | MYC_KO | NA | NA | NA | NA | NA | NA | NA |
| | | | MYCN_KO | 38 | 17.1171171 | 6.79863484 | 0.00148951 | 0.02764272 | 38 | 6.79863484 |
| | | | P3F_KO | 82 | -36.936937 | -27.270871 | 5.12E-27 | 1.94E-24 | -82 | -27.270871 |
| DAVICIONI_PAX_FOXO1_SIGNATURE_IN_ARMS_UP | M4991 | C2 | MYC_KO | 6 | 12.5 | 3.94887084 | 0.22568245 | 1 | 6 | 3.94887084 |
| | | | MYCN_KO | NA | NA | NA | NA | NA | NA | NA |
| | | | P3F_KO | 29 | -60.416667 | -50.444578 | 2.07E-17 | 1.79E-15 | -29 | -50.444578 |
| GRYDER_PAX3FOXO1_ENHANCERS_KO_DOWN | M227 | C2 | MYC_KO | NA | NA | NA | NA | NA | NA | NA |
| | | | MYCN_KO | 48 | -12.371134 | -1.7118986 | 0.16112683 | 1 | -48 | -1.7118986 |
| | | | P3F_KO | 103 | -26.546392 | -16.923471 | 9.79E-21 | 1.41E-18 | -103 | -16.923471 |
| GRYDER_PAX3FOXO1_TOP_ENHANCERS | M172 | C2 | MYC_KO | NA | NA | NA | NA | NA | NA | NA |
| | | | MYCN_KO | 47 | -11.380145 | -0.6889252 | 0.35169573 | 1 | -47 | -0.6889252 |
| | | | P3F_KO | 107 | -25.90799 | -16.298984 | 1.25E-20 | 1.72E-18 | -107 | -16.298984 |

**Supplementary Table 2: P3F-related gene sets affected in MYC, MYCN, and P3F knockdowns**

#### Supplementary Table 3

| Pathway_name | Pathway_Id | Categ | Group | Number_hits | Percent_gene_hits_per_pathway | Enrichment_score | pval | FDR | Net_number_hits | NES |
|---|---|---|---|---|---|---|---|---|---|---|
| BILD_MYC_ONCOGENIC_SIGNATURE | M2069 | C2 | MYC_KO | NA | NA | NA | NA | NA | NA | NA |
| | | | MYCN_KO | 69 | -37.5 | -27.206512 | 4.81E-22 | 8.10E-20 | -36 | -19.599443 |
| | | | P3F_KO | 24 | 13.0434783 | 2.4932943 | 0.16529783 | 0.49901852 | 2 | 0.68480028 |
| DANG_MYC_TARGETS_UP | M6506 | C2 | MYC_KO | 13 | -10.4 | -1.3926784 | 0.3375149 | 1 | -13 | -1.3926784 |
| | | | MYCN_KO | 54 | -43.2 | -32.830513 | 7.31E-21 | 1.01E-18 | -54 | -32.830513 |
| | | | P3F_KO | 32 | -25.6 | -15.588081 | 6.15E-07 | 1.68E-05 | -32 | -15.588081 |
| KIM_MYC_AMPLIFICATION_TARGETS_UP | M8445 | C2 | MYC_KO | 15 | -9.1463415 | -0.1260972 | 0.51777396 | 1 | -15 | -0.1260972 |
| | | | MYCN_KO | 62 | -37.804878 | -27.469181 | 3.87E-20 | 5.10E-18 | -62 | -27.469181 |
| | | | P3F_KO | 35 | -21.341463 | -11.321817 | 1.63E-05 | 3.66E-04 | -35 | -11.321817 |
| ACOSTA_PROLIFERATION_INDEPENDENT_MYC_TARGETS_UP | M2221 | C2 | MYC_KO | 7 | -9.8591549 | -0.8422058 | 0.46171716 | 1 | -7 | -0.8422058 |
| | | | MYCN_KO | 28 | -39.43662 | -28.894247 | 2.48E-10 | 1.50E-08 | -28 | -28.894247 |
| | | | P3F_KO | 17 | -23.943662 | -13.850442 | 6.06E-04 | 0.00998854 | -17 | -13.850442 |
| COLLER_MYC_TARGETS_UP | M5955 | C2 | MYC_KO | 3 | -13.636364 | -4.6229448 | 0.31878975 | 1 | -3 | -4.6229448 |
| | | | MYCN_KO | 18 | -81.818182 | -71.23501 | 1.49E-14 | 1.33E-12 | -18 | -71.23501 |
| | | | P3F_KO | 6 | -27.272727 | -17.128411 | 0.01961254 | 0.16483923 | -6 | -17.128411 |
| PID_MYC_ACTIV_PATHWAY | M66 | C2 | MYC_KO | 7 | -9.7222222 | -0.7045089 | 0.47669129 | 1 | -7 | -0.7045089 |
| | | | MYCN_KO | 28 | -38.888889 | -28.345623 | 3.65E-10 | 2.17E-08 | -28 | -28.345623 |
| | | | P3F_KO | 9 | -12.5 | -2.3381219 | 0.30889635 | 0.82254504 | -9 | -2.3381219 |
| SATOH_COLORECTAL_CANCER_MYC_UP | M42515 | C2 | MYC_KO | NA | NA | NA | NA | NA | NA | NA |
| | | | MYCN_KO | 43 | -53.75 | -43.326796 | 1.68E-21 | 2.68E-19 | -43 | -43.326796 |
| | | | P3F_KO | 23 | -28.75 | -18.699962 | 2.77E-06 | 6.72E-05 | -23 | -18.699962 |
| SCHLOSSER_MYC_TARGETS_AND_SERUM_RESPONSE_UP | M14278 | C2 | MYC_KO | NA | NA | NA | NA | NA | NA | NA |
| | | | MYCN_KO | 26 | -55.319149 | -44.78126 | 6.39E-14 | 5.24E-12 | -26 | -44.78126 |
| | | | P3F_KO | | NA | NA | NA | NA | NA | NA |

**Supplementary Table 3: Myc-related gene sets affected in MYC, MYCN, and P3F knockdowns**
